# Centrosomes control kinetochore-fiber plus-end dynamics via HURP to ensure symmetric divisions

**DOI:** 10.1101/557058

**Authors:** Damian Dudka, Nicolas Liaudet, Hélène Vassal, Patrick Meraldi

## Abstract

During mitosis centrosomes can affect the length of kinetochore-fibers (k-fibers) and the stability of kinetochore-microtubule attachments, implying that they regulate k-fiber dynamics. The exact cellular and molecular mechanisms by which centrosomes regulate k-fibers remain, however, unknown. Here, we created human non-cancerous cells with only one centrosome to investigate these mechanisms. Such cells formed highly asymmetric bipolar spindles that resulted in asymmetric cell divisions. K-fibers in acentrosomal spindles were shorter, more stable, had a reduced poleward microtubule flux at minus-ends, and more frequent pausing events at their plus-ends. This indicates that centrosomes regulate k-fiber dynamics both locally at minus-ends and far away at plus-ends. At the molecular level we find that the microtubule-stabilizing protein HURP is enriched on the k-fiber plus-ends in the acentrosomal spindles of cells with only one centrosome. HURP depletion rebalance k-fiber stability and dynamics in such cells, and improved spindle and cell division symmetry. Our data further indicate that HURP accumulates on k-fibers inversely proportionally to half-spindle length. We propose that centrosomes regulate k-fiber plus-ends indirectly via length-dependent accumulation of HURP. Thus by ensuring equal k-fiber length, centrosomes promote HURP symmetry, reinforcing the symmetry of the mitotic spindle and of cell division.

## INTRODUCTION

The bipolar mitotic spindle is a transient microtubule-based structure that ensures faithful chromosome segregation in all eukaryotic cell divisions. In most metazoans, the main mitotic microtubule-organizing centers are the centrosomes (Prosser and Pelletier, 2017). At mitotic onset, centrosome-nucleated microtubules capture chromosomes via kinetochores and align them at the spindle equator to form a metaphase plate (Kapoor, 2017). Centrosomes at each spindle pole are composed of two microtubule-based centrioles and pericentriolar material that nucleates microtubules (Wu and Akhmanova, 2017). Centrosomes and the associated centrioles duplicate once per cell cycle under the control of the Plk4 kinase (Loncarek and Bettencourt-Dias, 2018). Despite their well-described role in promoting bipolar spindle assembly, centrosomes are absent in higher plants, planarians, and during the first divisions of mouse embryos (Azimzadeh et al., 2012; Yi and Goshima, 2018; Zamboni, 1970). They are also dispensable for bipolar spindle formation in *Drosophila melanogaster* (Basto et al., 2006), *Xenopus laevis* egg extracts (Heald et al., 1996) and mammalian somatic cells (Khodjakov et al., 2000). Centrosomes are, however, required for mitotic bipolar spindle assembly in *Caenorhabditis elegans* (Delattre et al., 2004) and sea urchin (Sluder and Rieder, 1985). Centrosomes also ensure faithful chromosome segregation, since somatic centrosome-free fly, chicken or human cells often mis-segregate chromosomes, despite achieving bipolar spindles (Buffin et al., 2007; Sir et al., 2013; Wong et al., 2015).

Centrosomes have been proposed to act in a dominant manner on spindle formation and spindle shape. The presence of a single centrosome results in the formation of monopolar spindles in *C. elegans*, sea urchins and *X. leavis* egg extracts (Heald et al., 1997; Mazia et al., 1960; O’Connell et al., 2001) and a mixed population of bipolar and monopolar spindles in human cancer cells (Leidel et al., 2005). Removing a centriole in one centrosome leads to asymmetric spindles in the green alga *Chlamydomonas reinhardtii* (Keller et al., 2010), *C. elegans* (Greenan et al., 2010) and human cells (Tan et al., 2015). Finally, naturally occurring unequal amount of the pericentriolar material results in asymmetric spindles in the annelid *Helobdella robusta* zygotes (Ren and Weisblat, 2006) and mollusk embryos (Dan and Inoué, 1987).

One mechanism by which centrosomes regulate the mitotic spindle is the control of kinetochore-microtubule dynamics. Kinetochore-microtubules form bundles (k-fibers) that are intrinsically polarized structures with dynamic minus-ends embedded at centrosomes and plus-ends attached to kinetochores (Akhmanova and Steinmetz, 2015). K-fibers dynamics have to be tightly regulated since they contribute to spindle assembly, chromosome attachment and congression, kinetochore oscillations on the metaphase plate, half-spindle length, correction of erroneous kinetochore-microtubule attachments, and finally the synchronicity of anaphase movements (Amaro et al., 2010; Cimini et al., 2004; Dudka et al., 2018; Jaqaman et al., 2010; Stumpff et al., 2008; Tan et al., 2015; Toso et al., 2009; Vladimirou et al., 2013; Wordeman et al., 2007).

Centrosomes regulate k-fiber minus-end dynamics via microtubule depolymerases of the kinesin-13 family such as Kif2a and MCAK, and microtubule-severing enzymes, such as Katanin (Ganem et al., 2005; Jiang et al., 2017). Together these enzymes generate poleward microtubule flux (Ganem et al., 2005; Jiang et al., 2017; Mitchison, 1989). Surprisingly, centrosomes also influence k-fiber plus-end dynamics even tough the kinetochore-microtubule interface is located far away. We have previously shown that single centriole ablation influence half-spindle lengths and delays maturation of the kinetochore-microtubule attachments (Tan et al., 2015), and that centrosome age sets the relative stability of the kinetochore-microtubule attachments (Gasic et al., 2015). The mechanisms by which centrosomes influence k-fiber plus-end dynamics are, however, unknown.

K-fiber plus-end dynamics are set by kinetochores themselves, microtubule-depolymerases such as Kif2a, MCAK or Kif18A (Amaro et al., 2010; Ganem et al., 2005; Mayr et al., 2007; Wordeman et al., 2007) and plus-end binding proteins, such as CLASP, ch-TOG or HURP (Barr and Gergely, 2008; Maiato et al., 2003; Sillje et al., 2006). Of particular interest is the k-fiber stabilizing HURP-protein, since it is specifically enriched on a k-fiber section proximal to kinetochores; the mechanisms governing its function and localization are, however, unclear (Koffa et al., 2006; Sillje et al., 2006; Wong and Fang, 2006; Wong et al., 2008; Wu et al., 2013). Finally, k-fiber dynamics are also affected by k-fiber length, which is regulated by microtubule motors such as HSET, Kid, and Kif15 (Cai et al., 2009; Mayr et al., 2007; Sturgill and Ohi, 2013; Tokai-Nishizumi et al., 2005).

Here, we investigated the role of centrosomes in k-fiber regulation by creating human cells with bipolar spindles containing a single centrosome. This allowed us to perform a direct side-by-side comparison of k-fiber dynamics in half spindles with and without centrosomes. We find that acentrosomal k-fibers are more stable, more prone to pausing events, and shorter, resulting in asymmetric spindles and asymmetric cell divisions. These differences partially depend on HURP, which we find to be enriched on the plus-ends of the acentrosomal k-fibers. HURP asymmetry results from its ability to accumulate on shorter k-fibers, whose length is set by the presence or absence of centrosomes. Overall, our results lead us to propose that centrosomes regulate plus-end k-fiber dynamics by indirectly controlling HURP localization to promote symmetric cell divisions.

## RESULTS

### Centrosome ablation leads to asymmetric spindles and asymmetric divisions

To study how centrosomes affect the mitotic spindle we removed them by blocking centrosome duplication with the Plk4 inhibitor centrinone (Wong et al., 2015). Human retina pigment epithelial cells immortalized with telomerase (hTert-RPE1) expressing the microtubule and spindle pole marker EB3-GFP and the chromosome marker H2B-mCherry, were treated for 32 hours with 300 nM centrinone. This led to formation of monastral bipolar spindles with only one EB3-GFP aster (Fig. 1A). Immunofluorescence staining against the centriole marker centrin-1 indicated that this aster contained one centrosome with a single centriole, and no centriole at the opposite spindle pole (Fig. 1A). These centrinone-treated cells were called 1:0 cells as opposed to untreated 2:2 cells (with two centrioles at each pole). Live cell imaging indicated that 1:0 hTert-RPE1 EB3-GFP/H2B-mCherry cells first assembled monopolar spindles before forming asymmetric bipolar spindles; the longer half-spindle emanated from the centrosomal spindle pole (Fig. 1B; Movies S1 and S2). The monopolar to bipolar transition prolonged the overall mitotic timing in 1:0 cells (36 +/− 3 min 95 % CI vs. 15 +/− 0 min in 2:2 cells; Fig. 1C), but did not affect the time between chromosome congression and anaphase onset (Fig. S1A). 1:0 cells also had a higher incidence of lagging chromosomes in anaphase, indicating a higher rate of chromosome mis-segregation (13.1 % vs. 2.2 % in 2:2 cells; Fig. 1D). We conclude that the presence of a single centrosome delays but does not prevent bipolar spindle formation, and that it affects spindle symmetry and chromosome segregation.

**Figure 1.**
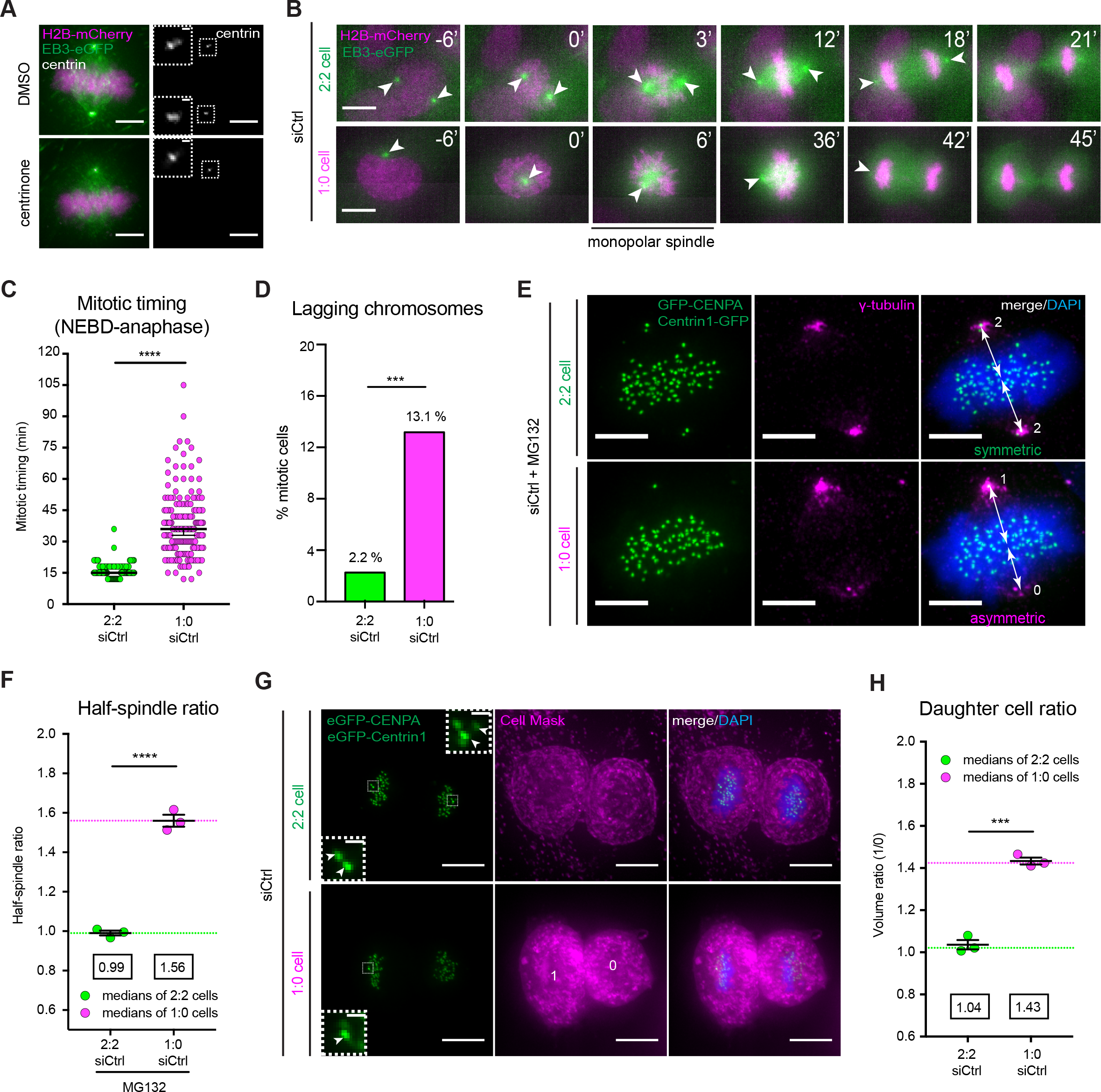
Centrosome ablation leads to asymmetric spindles and asymmetric divisions. (**A**) Immunofluorescence images of hTert-RPE1 EB3-GFP/H2B-mCherry cells treated with DMSO or 300 nM of the Plk4 inhibitor centrinone for 32 h and stained for centrin1 (white). Insets show the centrioles in each spindle pole. Scale bars = 5 μm (**B**) Time-lapse images of control-depleted hTert-RPE1 EB3-GFP/H2B-mCherry cells treated for 32 h with either DMSO (2:2 cell) or 300 nM centrinone (1:0 cell). White arrowheads point at centrosomes. Scale bars = 5 μm; (see Supplementary Movie 1 and 2). (**C**) Mitotic timing measured from nuclear envelope breakdown (NEBD) until anaphase onset in control-depleted hTert-RPE1 EB3-GFP/H2B-mCherry 2:2 or 1:1 cells; N = 4, n = 143-182 cells; error bars represent median and 95% CI; Mann-Whitney test; p < 0.0001. (**D**) Quantification of lagging chromosomes in control-depleted 2:2 and 1:0 hTert-RPE1 EB3-GFP/H2B-mCherry cells; N = 4, n = 99-134 cells; Fisher’s exact test with 99% CI; p < 0.001. (**E**) Immunofluorescence images of control-depleted 2:2 and 1:0 hTert-RPE1 Centrin1-GFP/GFP-CENPA cells blocked in metaphase prior fixation, and stained with γ-tubulin antibodies and DAPI. Scale bars = 5 μm. White arrows indicate the distance between the poles and the center of the metaphase plate. (**F**) Quantification of the half-spindle ratio (in 3D) in control-depleted 2:2 and 1:0 hTert-RPE1 Centrin1-GFP/GFP-CENPA cells blocked in metaphase prior fixation; dashed lines show the mean half-spindle ratio (green – 2:2 cells; magenta – 1:0 cells); N = 3, n = 150-169 cells; error bars represent s.e.m.; two-tailed unpaired t-test; p < 0.0001. (**G**) Immunofluorescence images of control-depleted 2:2 and 1:0 hTert-RPE1 Centrin1-GFP/GFP-CENPA cells in cytokinesis, fixed and stained with Cell Mask and DAPI. Scale bars = 5 μm. Inserts show centrioles. Scale bars = 0.5 μm. (**H**) Quantification of the mean daughter cell ratio (see Methods section); N = 3, n = 91-99 cells; error bars indicate s.e.m.; two-tailed unpaired t-test; p = 0.0001. *** Indicates p < 0.001, **** indicates p < 0.0001.

To quantify this asymmetry over time, we synchronized hTert-RPE1 EB3-GFP/H2B-mCherry cells with the Cdk1 inhibitor RO-3306 and released them into medium containing the proteasome inhibitor MG132 to maintain them in metaphase. The ratio of the two half-spindles started approximately at 1.7 (centrosomal/acentrosomal), before reaching a steady state of roughly 1.5 after 15 min (Fig. S1B). This showed that centrosomes regulate k-fiber length. Thanks to the presence of spindle pole marker γ-tubulin at both poles, we precisely quantified by immunofluorescence in 3D the half-spindle ratios in MG312-arrested 1:0 hTert-RPE1 Centrin1-GFP (centriole marker)/GFP-CENPA (kinetochore marker) cells (Fig. 1E and Fig. S1C). 1:0 cells had a median half-spindle ratio of 1.56 ± 0.03 s.e.m. versus 0.99 ± 0.01 s.e.m. in 2:2 cells (Fig. 1F and Fig. S1D). This half-spindle ratio was independent of Plk4 inhibition, since depletion of the essential centrosome duplication component Sas6, or depletion of Sas6 combined with acute Plk4 inhibition resulted in similar ratios as Plk4 inhibition alone (Fig. S1E and F).

Since spindle symmetry contributes to the symmetry of cell divisions (Tan et al., 2015), we further stained 2:2 and 1:0 hTert-RPE1 Centrin1-GFP/GFP-CENPA cells with the plasma membrane marker Cell Mask to quantify the volume of the two daughter cells in telophase. While 2:2 cells divided symmetrically (ratio of 1.04 ± 0.02), 1:0 cells showed asymmetric divisions with the centrosome associated to the larger daughter cell (1.43 ± 0.02, Fig. 1G and H). Overall, our results show that loss of a single centrosome does not prevent bipolar spindle assembly in human cells, but that it leads to severe spindle asymmetry and asymmetric cell divisions, a phenotype that has been previously linked to differential k-fiber regulation (Tan et al., 2015).

### Centrosome ablation affects both minus and plus-end k-fiber dynamics

The asymmetric spindles of 1:0 cells provided a unique model to test side-by-side how the presence or absence of centrosomes affects k-fibers. To evaluate k-fiber stability, we cold-treated MG132-arrested hTert-RPE1 Centrin1-GFP/GFP-CENPA 1:0 cells for 5 min to eliminate short-lived non-kinetochore microtubules and induce k-fiber depolymerization. Each half-spindle was classified as: intact half-spindle structures (class 1), half-spindles with several k-fibers missing (class 2), or half-spindles with most k-fibers missing (class 3; Fig. 2A). K-fibers in acentrosomal half-spindles tended to be more intact than the centrosomal ones, suggesting higher stability (Fig. 2B). Nevertheless, since this trend was not significant (p = 0.0514 in Chi square test), we aimed to validate or refute it with a second, independent assay. We labeled MG132-arrested hTert-RPE1 Centrin1-GFP/GFP-CENPA 1:0 cells with the live dye SiR-tubulin (Lukinavicius et al., 2014), allowing us to visualize both k-fibers and centrioles, before treating them with a 200 ng/ml spike of the microtubule-depolymerizing drug nocodazole (Fig. 2C). We found that k-fibers in centrosomal half-spindles lost their integrity 3 min earlier than in acentrosomal half-spindles (5min 38s ± 28 s versus 8min 18s ± 38 s; p = 0.003 in paired t-test; Fig. 2D). This confirmed that in 1:0 cells the k-fibers in the acentrosomal half-spindle are more stable.

**Figure 2.**
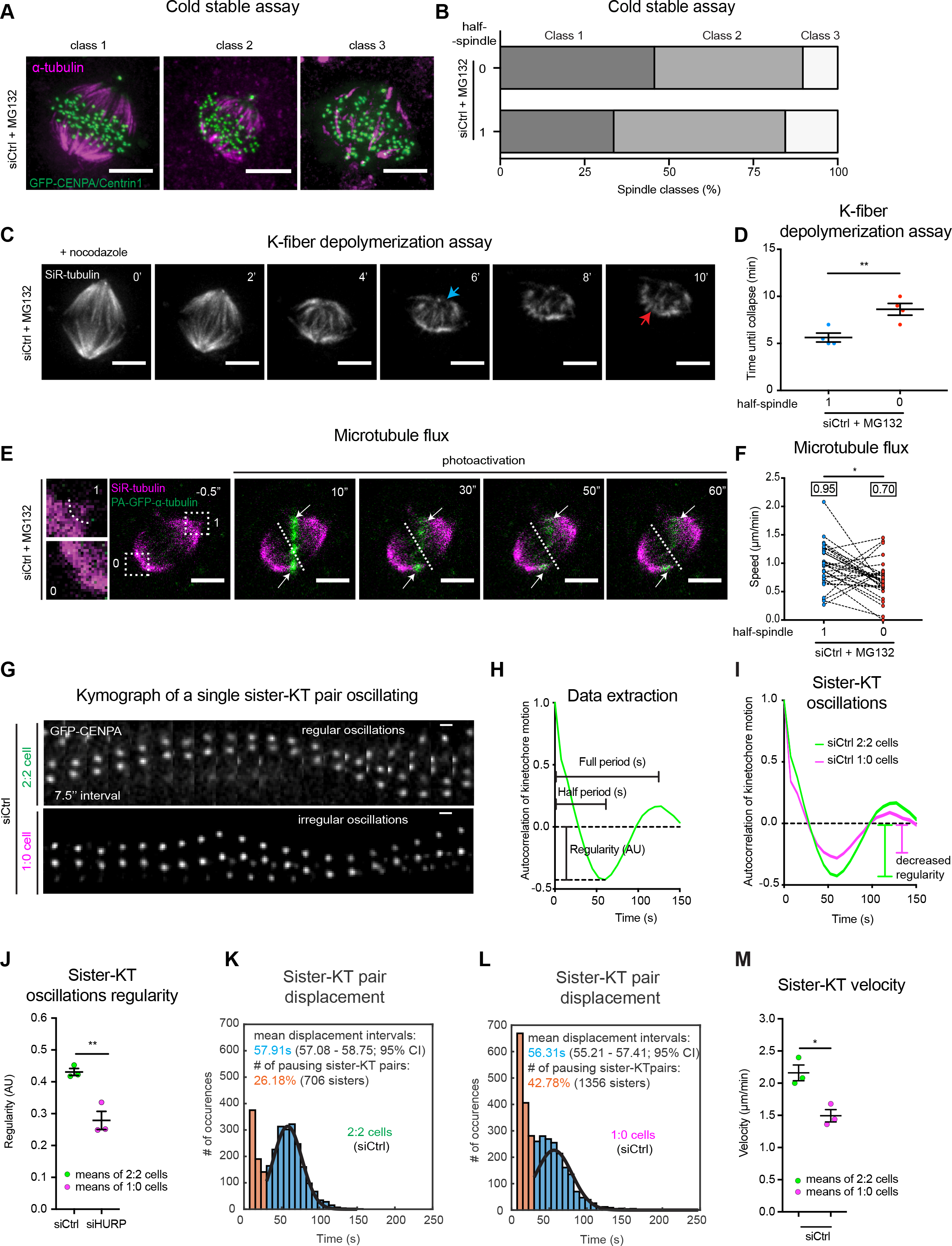
Centrosome ablation affects both minus and plus-end k-fiber dynamics. (**A**) Immunofluorescence images of control-depleted 1:0 hTert-RPE1 Centrin1-GFP/GFP-CENPA cells blocked in metaphase and treated with cold for 5 min fixed and stained with α-tubulin antibodies. The classes illustrating cold-stability are explained in the main text. Scale bars = 5 μm (**B**) Quantification of the cold stable classes; N = 4, n = 112-116 cells. (**C**) Time-lapse images of the k-fiber depolymerization assay in control-depleted 1:0 hTert-RPE1 Centrin1-GFP/GFP-CENPA cells stained with SiR-tubulin and blocked in metaphase. Scale bars = 5 μm. Blue arrow indicates the collapse of “1” half-spindle, red arrow indicates the collapse in “0” half-spindle. (**D**) Quantification of the k-fiber depolymerization assay; N = 4, n = 109 cells; error bars represent mean and s.e.m.; two-tailed unpaired t-test; p = 0.0034. (**E**) Time-lapse images of photo-activation in control-depleted 1:0 hTert-RPE1 PA-GFP-α-tubulin cells stained with SiR-tubulin and blocked in metaphase. Scale bars = 5 μm. Inserts demonstrate the spindle pole morphology used to distinguish “1” and “0” poles. White arrows follow the photo-activated regions of k-fibers. Dashed lines indicate an approximate metaphase plate position. (**F**) Quantification of the microtubule flux in control-depleted 1:0 hTert-RPE1 PA-GFP-α-tubulin cells; N = 3, n = 31 cells; mean values are indicated in boxes; two-tailed paired t-test; p = 0.0117. (**G)** Kymograph of a representative single, oscillating sister-KT pair from a metaphase control-depleted 2:2 and 1:0 hTert-RPE1 Centrin1-GFP/GFP-CENPA cells. Scale bars = 1 μm; (see Supplementary Movies 3 and 4). **H**) Representation of data extraction from autocorrelation curves of sister-KT pair oscillations. (**I**) Cumulative autocorrelation curves representing sister-KT oscillations in control-depleted 2:2 (green curve) and 1:0 (magenta curve) hTert-RPE1 Centrin1-GFP/GFP-CENPA cells. Vertical bars represent the regularity of the oscillations, higher in 2:2 cells (green bar) than in 1:0 cells (magenta bar). N = 3, n = 36 cells; n_kt_ = 1368-1380 pairs. Line thickness indicates s.d. (**J**) Quantification of the regularity of sister-KT oscillations represented in H. Error bars represent means and s.e.m.; N = 3, n = 36 cells; n_kt_ = 1368-1380 pairs. (**K** and **L**) Histograms of sister-KT pair displacement in control-depleted 2:2 (K) and 1:0 (L) hTert-RPE1 Centrin1-GFP/GFP-CENPA cells; orange – pausing pairs; blue – mean displacement intervals of dynamic pairs, black line – normal distribution fit. N = 3, n = 36 cells; n_kt_ = 1368-1380 pairs. (**M**) Sister-KT velocity in control-depleted 2:2 and 1:0 cells. Each line shows how many sister-KT pairs were detected moving by a given distance from one frame to another. The velocity was quantified as a mean displacement of all sister-KT pairs, multiplied by the number of frames within 1 minute of a movie (= 8). N = 3, n = 36 cells; n_kt_ = 1368-1380 pairs. * Indicates p < 0.05, ** indicates p < 0.01.

K-fiber dynamics are determined both at the minus- and the plus-ends (Akhmanova and Steinmetz, 2015). To verify the impact of centrosome ablation on k-fiber minus-ends we measured microtubule flux, a poleward conveyer belt of tubulin dimers within k-fibers, powered by the k-fiber depolymerization at spindle poles (Ganem et al., 2005; Mitchison, 1989). We quantified flux rates in 1:0 cells using hTert-RPE1 cells expressing photoactivatable-GFP-α-tubulin and stained them with 50 nM SiR-tubulin to visualize spindle poles and centrioles. To simultaneously measure the flux in both half-spindles, we photoactivated k-fibers in MG132-arrested 1:0 cells across the metaphase plate (Fig. 2E). We found a slower median flux rate in the acentrosomal k-fibers (0.70 ± 0.33 μm/min s.d. versus 0.95 ± 0.38 μm/min s.d. in centrosomal half-spindles; Fig. 2F). Importantly, the rate of flux in the centrosomal half-spindle was identical to the value previously measured in 2:2 cells not treated with SiR-tubulin (Toso et al., 2009), indicating that 50nM SiR-tubulin did not interfere with flux. We conclude that centrosome loss reduces k-fiber depolymerization at spindle poles.

As a read-out for k-fiber plus-end dynamics, we used a kinetochore tracking software that detects sister-kinetochore pairs and extracts the parameters describing their movements (Jaqaman et al., 2010) (Fig. 2G). In metaphase, sister-kinetochores oscillate along the spindle axis, reflecting the dynamic instability of k-fiber plus-ends (Amaro et al., 2010; Jaqaman et al., 2010). By applying an auto-correlation analysis one obtains a sinusoidal curve, in which the position of the first minimum indicates the mean half-period of the oscillations, and the depth of the minimum indicates the regularity of the oscillations (Olziersky et al., 2018) (Fig. 2H). We found that sister-kinetochores oscillate less regularly in 1:0 cells than in 2:2 cells (Fig. 2I and J; Movies S3 and S4). When we looked at the duration of individual kinetochore trajectories, we found two populations: a first population that switched direction on average every 55 seconds (blue), representing kinetochores bound to persistently growing or shrinking k-fibers (Fig. 2K and L); and a second population (orange) kinetochores displaying the signature of pausing plus-ends: apparent jittering movements within the 7.5s time resolution reflecting the uncertainty in positioning measurements of our assay (Fig. 2K and L). While the period of the oscillating kinetochores was the same in 2:2 and 1:0 cells (Fig. 2I), the proportion of kinetochores bound to pausing microtubules was much higher in 1:0 cells (43% vs. 26%; Fig. 2K and L). Moreover, the average kinetochore velocity was reduced when compared to 2:2 cells (Fig. 2M). This confirmed that loss of centrosomes changes k-fiber dynamics not only at minus-, but also at plus-ends.

### HURP depletion increases spindle and cell division symmetry in 1:0 cells

Our previous work indicated that spindle asymmetry depends on an imbalance in k-fiber dynamics (Tan et al., 2015). We reasoned that screening for depletions of k-fiber regulators that accentuate or reduce spindle asymmetry in 1:0 cells could identify candidates through which centrosomes control k-fiber dynamics. When we tested how depletion of ch-TOG, CLASP-1, Kid, Kif2a, Kif-15, HSET or HURP affected half-spindle ratios in 1:0 cells, we only found a reduction after HURP or ch-Tog depletion (Fig. S2A and S2B) – Kif18A depletion could not be evaluated, since it disrupted spindle assembly in 1:0 cells. Depletion of ch-TOG in addition reduced spindle length suggesting a severe effect on spindle function (Fig. S2B); in contrast, HURP depletion only had a minor effect on spindle length (Fig. S2B). Since HURP depletion gave a more specific phenotype, we focused on the potential link between centrosomes, HURP function, and k-fiber plus-end dynamics.

To validate HURP as hit, we confirmed that HURP depletion significantly increased spindle symmetry in 1:0 cells in multiple replicates and when using two different siRNAs, without affecting spindle ratio in 2:2 cells (Fig. 3A-C, and Fig. S2C and D). Immunofluorescence also confirmed that both HURP siRNA oligonucleotides were efficient at abolishing the well-documented k-fiber plus-end localization of HURP ((Sillje et al., 2006); Fig. 3D). We next tested if the partial rescue of spindle asymmetry seen after HURP depletion would also rescue the asymmetric cell divisions. HURP depletion with either siRNA did not affect the symmetric divisions in 2:2 cells (daughter cell ratio of 0.99 +/− 0.03 s.e.m. for siHURP vs. 1.04 +/− 0.02 s.e.m. in siCtrl cells) but partially restored the symmetry in 1:0 cells (1.24 +/− 0.02 s.e.m. vs. 1.43 +/− 0.02 s.e.m. in siCtrl cells; Fig. 3E-G). In contrast, when we monitored spindle assembly and chromosome segregation by live cell imaging, we found that HURP-depleted 1:0 cells still transited from a monopolar spindle to bipolar spindle configuration with a mild delay compared to control-depleted 1:0 cells (Fig. S2E-I), and that their chromosome segregation error rate was only mildly lower than in control-depleted 1:0 cells (Fig. 3H). We conclude that HURP depletion increases spindle and cell division symmetry in 1:0 cells.

**Figure 3.**
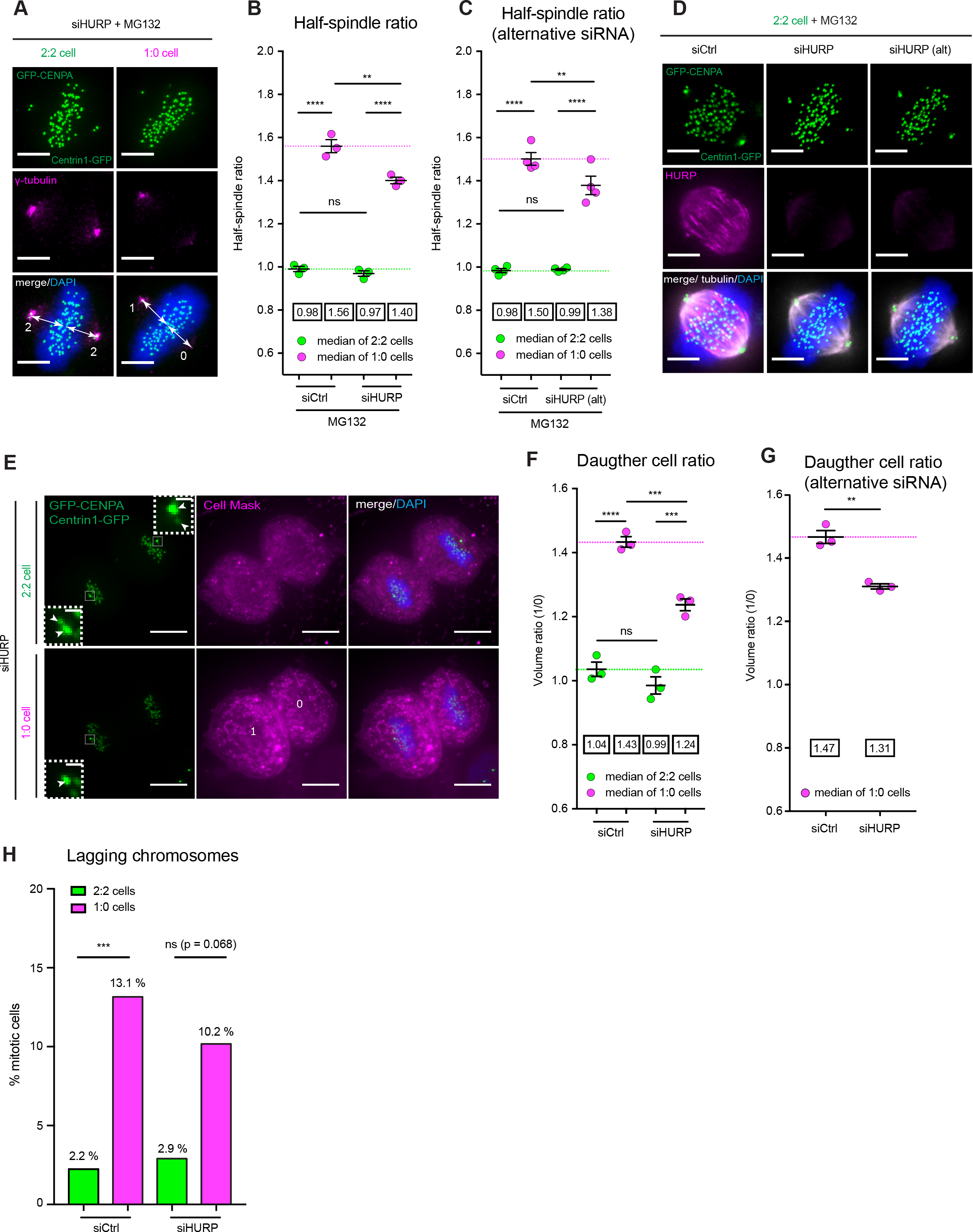
HURP depletion increases spindle and cell division symmetry in 1:0 cells. (**A**) Immunofluorescence images of 2:2 and 1:0 hTert-RPE1 Centrin1-GFP/GFP-CENPA cells metaphase cells depleted of HURP, stained with γ-tubulin antibodies and DAPI. Scale bars = 5 μm. White arrows point at the distance between the poles and the center of the metaphase plate. (**B** and **C**) Quantification of the half-spindle ratio (in 3D) in 2:2 and 1:0 hTert-RPE1 Centrin1-GFP/GFP-CENPA cells blocked in metaphase after control and HURP depletion (B) or after treatment with an alternative siHURP oligonucleotide (C); dashed lines show the mean half-spindle ratio for control-depleted 2:2 (green) and 1:0 (magenta) cells; N = 3-4, n = 150-181 cells; error bars represent mean and s.e.m.; two-way ANOVA. 2:2 siCtrl vs. 1:0 siCtrl p < 0.0001; 2:2 siHURP vs. 1:0 siHURP p < 0.0001; 1:0 siCtrl vs. 1:0 siHURP p = 0.006; (B) and 2:2 siCtrl vs. 1:0 siCtrl p < 0.0001; 2:2 siHURP(alt) vs. 1:0 siHURP(alt) p < 0.0001; 1:0 siCtrl vs. 1:0 siHURP(alt) p = 0.0075 (C) in two-way ANOVA. Note that the control-depleted cells are the same than those reported in Fig. 1. (**D**) Immunofluorescence images of control- and HURP-depleted 2:2 hTert-RPE1 Centrin1-GFP/GFP-CENPA metaphase cells, stained with γ-tubulin antibodies and DAPI. Scale bars = 5 μm. siHURP(alt) represents an alternative siHURP oligonucleotide used. White arrows point at the distance between the poles and the center of the metaphase plate. (**E**) Immunofluorescence images of control- and HURP-depleted 2:2 and 1:0 hTert-RPE1 Centrin1-GFP/GFP-CENPA cells in cytokinesis, stained with Cell Mask and DAPI. Scale bars = 5 μm. Inserts show centrioles. Scale bars = 0.5 μm. (**F** and **G**) Quantification of the daughter cell ratio (see Methods section) after HURP depletion (F) or treatment with an alternative siHURP oligonucleotide (G); N = 3, n = 91-138 cells; mean values are indicated in boxes; error bars indicate mean and s.e.m.; 2:2 siCtrl vs. 1:0 siCtrl p < 0.0001; 2:2 siHURP vs. 1:0 siHURP p = 0.0005; 1:0 siCtrl vs. 1:0 siHURP p = 0.0018 in two-way ANOVA (F) and 1:0 siCtrl vs. 1:0 siHURP(alt) p = 0.0021 (G) in unpaired t-test. HURP-depleted cells were compared with the control-depleted cells reported in Fig. 1. (**H**) Quantification of lagging chromosomes in control- and HURP-depleted 2:2 and 1:0 hTert-RPE1 EB3/H2B-mCherry cells; N = 4, n = 59-138 cells; Fisher’s exact test 99% CI; 2:2 siCtrl vs. 1:0 siCtrl p < 0.001; 2:2 siHURP vs. 1:0 siHURP p = 0.0684; 2:2 siCtrl vs. 2:2 siHURP p > 0.9999; 1:0 siCtrl vs. 1:0 siHURP p = 0.6322. ** Indicates p < 0.01, *** indicates p < 0.001, **** indicates p < 0.0001, ns – not significant.

### HURP depletion rescues plus-end k-fiber dynamics

HURP stabilizes k-fibers (Sillje et al., 2006), raising the possibility that its depletion may promote equal k-fiber lengths in 1:0 cells by regulating k-fiber dynamics. In the cold stable assay, HURP depletion abolished the difference in k-fiber stability between the centrosomal and the acentrosomal half-spindles in 1:0 cells – moreover it reduced the overall stability of the spindle (Fig. 4A and B). A similar result could be seen after HURP depletion in the nocodazole-depolymerization assay: the overall intensity of the spindle decreased faster when compared to control depletion (Fig. 4C) and the difference between k-fiber stability in the centrosomal and acentrosomal half-spindles was to reduced to 1.5 min (Fig. 4D and E). We conclude that HURP depletion partially suppresses the difference in k-fiber stability in 1:0 cells.

**Figure 4.**
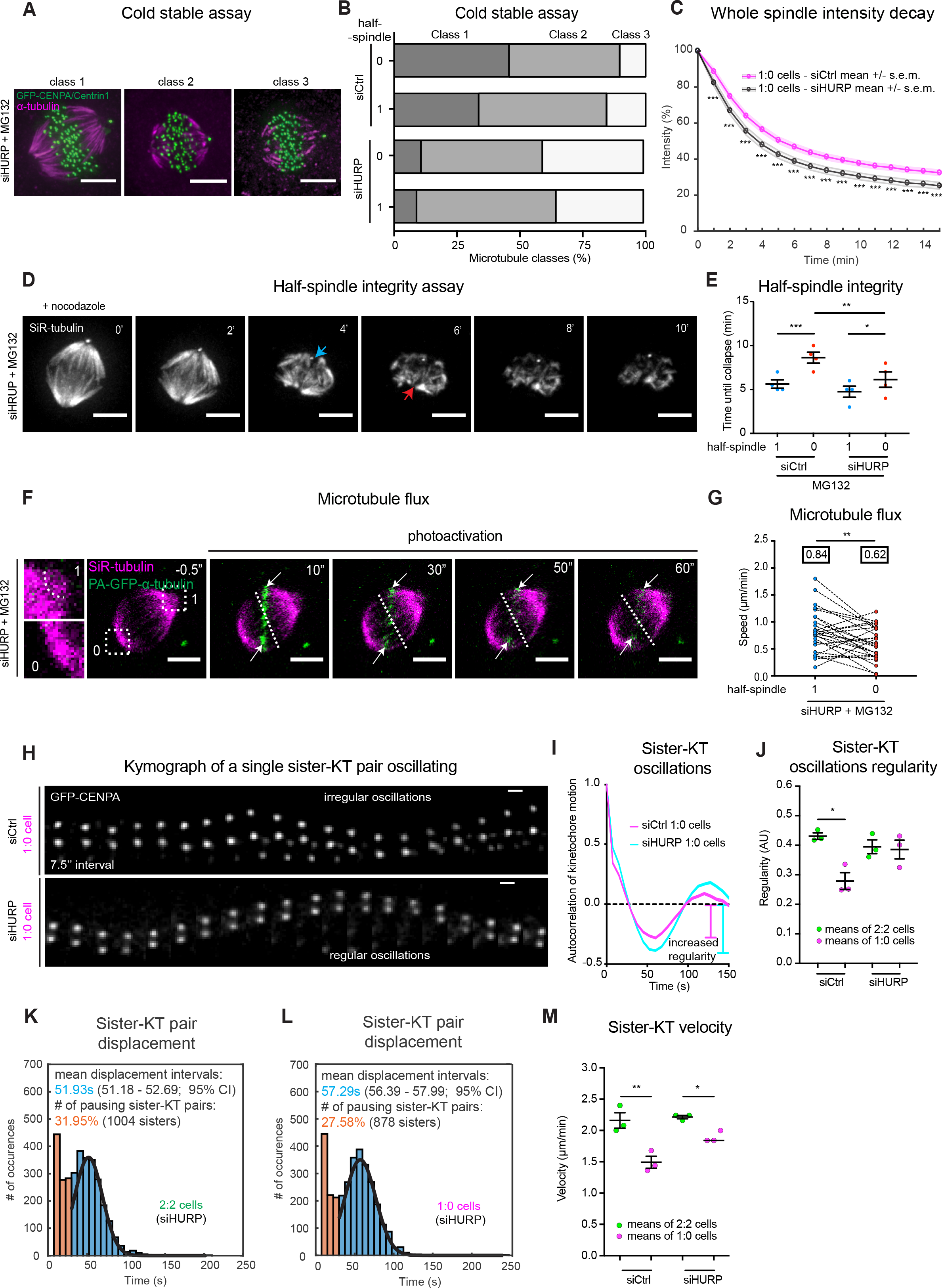
HURP depletion rescues plus-end k-fiber dynamics. (**A**) Immunofluorescence images of HURP-depleted 1:0 hTert-RPE1 Centrin1-GFP/GFP-CENPA cells blocked in metaphase, treated with ice-cold medium for 5 min; and stained with α-tubulin antibodies. Scale bars = 5 μm. (**B**) Quantification of the cold stable assay; N = 4, n = 112-116 cells. HURP-depleted cells were compared with the control-depleted cells reported in Fig. 2. (**C**) Quantification of the microtubule depolymerization assay in the whole spindle in control- (magenta) and HURP- (black) depleted 1:0 hTert-RPE1 Centrin1-GFP/GFP-CENPA cells stained with SiR-tubulin, blocked in metaphase and treated with 200 ng/ml of nocodazole; N = 4, n = 108-109 cells; error bars represent mean and s.e.m.; repeated measures ANOVA test; p < 0.001. (**D**) Time-lapse images of the microtubule depolymerization assay in HURP-depleted 1:0 hTert-RPE1 Centrin1-GFP/GFP-CENPA cells stained with SiR-tubulin and blocked in metaphase. Scale bars = 5 μm. Blue arrow indicates the collapse of “1” half-spindle, red arrow indicates the collapse in “0” half-spindle. (**E**) Quantification of the microtubule depolymerization assay; N = 4, n = 108-109 cells; error bars represent mean and s.e.m.; 1 vs. 0 siCtrl half-spindle p = 0.0003; 1 vs. 0 siHURP half-spindle p = 0.0426; siCtrl vs. siHURP 0 half-spindle p = 0.0011 in two-way ANOVA. HURP-depleted cells were compared with the control-depleted cells reported in Fig. 2. (**F**) Time-lapse images of photo-activation in HURP-depleted 1:0 hTert-RPE1 PA-GFP-α-tubulin cells stained with SiR-tubulin and blocked in metaphase. Scale bars = 5 μm. Inserts demonstrate the spindle pole morphology used to distinguish “1” and “0” poles. White arrows follow the photo-activated regions of kinetochore-microtubules. Dashed lines indicate the approximate metaphase plate position. (**G**) Quantification of the microtubule flux in control-depleted 1:0 hTert-RPE1 PA-GFP-α-tubulin cells; N = 3, n = 29 cells; mean values are indicated in boxes; two-tailed paired t-test; p = 0.0051. (**H**) Kymograph of a representative single, oscillating sister-KT pair from metaphase control- and HURP-depleted 1:0 hTert-RPE1 Centrin1-GFP/GFP-CENPA cells. The kymograph for 1:0 control-depleted pair was already shown in Fig. 2D. Scale bars = 1 μm; (see Supplementary Movies 7 and 8). (**I**) Cumulative autocorrelation curves representing sister-KT oscillations in control- (magenta) and HURP- (cyan) depleted 1:0 hTert-RPE1 Centrin1-GFP/GFP-CENPA cells. Line thickness indicates s.d. Vertical bars represent the regularity of the oscillations. N = 3, n = 36 cells; n_kt_ = 1289-1457 pairs. **J**) Quantification of the regularity of sister-KT oscillations of control- and HURP-depleted, 2:2 and 1:0 hTert-RPE1 Centrin1-GFP/GFP-CENPA cells. Error bars represent means and s.e.m.; N = 3, n = 36 cells; n_kt_ = 1289-1457 pairs. The data from control-depleted 2:2 and 1:0 cells were already shown in Fig. 2J. (**K** and **L**) Histograms of sister-KT pair displacement in 2:2 HURP-depleted (K) and 1:0 HURP-depleted (L) hTert-RPE1 Centrin1-GFP/GFP-CENPA cells; orange – pausing pairs; blue – mean displacement intervals, black line – normal distribution fit. N = 3, n = 36 cells; n_kt_ = 1289-1457 pairs. (**M**) Sister-KT velocity in control- and HURP-depleted 1:0 cells. Each line shows how many sister-KT pairs were detected moving by a given distance from one frame to another. Error bars represent means and s.e.m.; N = 3, n = 36 cells; n_kt_ = 1289-1457 pairs. * Indicates p < 0.05, ** indicates p < 0.01, *** indicates p < 0.001.

When we looked how HURP depletion affected k-fiber dynamics at minus ends, we found that it did not alleviate the slower flux rates in the acentrosomal half-spindles of 1:0 cells (Fig. 4F and G). At k-fiber plus-ends in contrast, HURP depletion restored the regularity of sister-kinetochore oscillations in 1:0 cells to levels equivalent to 2:2 cells (Fig. 4H-J; Fig. S3A and B; Movie S7 and S8). It also suppressed the pausing events typical for 1:0 cells (Fig. 4K and L) and improved kinetochore velocity (Fig. 4M). Moreover, HURP depletion reduced the mean inter-kinetochore distances in both 2:2 and 1:0 cells (Fig. S3C), consistent with previous studies (Wong and Fang, 2006). Overall our results implied that centrosome loss affects k-fiber plus-end dynamics via HURP.

### HURP is enriched on the k-fibers of acentrosomal half-spindles in 1:0 cells

To better understand how centrosomes might affect HURP function, we studied its localization by immunofluorescence staining. This revealed a striking difference in HURP localization in 1:0 cells. HURP still localized to k-fiber plus-ends, as in 2:2 cells, but it showed a strong enrichment on the k-fibers of the acentrosomal half-spindle (Fig. 5A). We quantified this asymmetry using an in-house MATLAB-based 3D line-profiling tool, which allows an unbiased and rapid analysis of hundreds of mitotic spindles (Fig. 5B and Material and methods). Our quantification showed that acentrosomal half-spindles had slightly higher tubulin levels only in the proximity of the poles (by 10%) and were strongly enriched for HURP (by 35%) close to the metaphase plate (Fig. 5C). Quantification with an automated kinetochore-microtubule intensity macro (Dudka et al., 2018) confirmed that HURP was enriched at k-fiber plus-ends in the acentrosomal half-spindle of 1:0 cells, even when normalized to tubulin (Fig. S4C and D).

**Figure 5.**
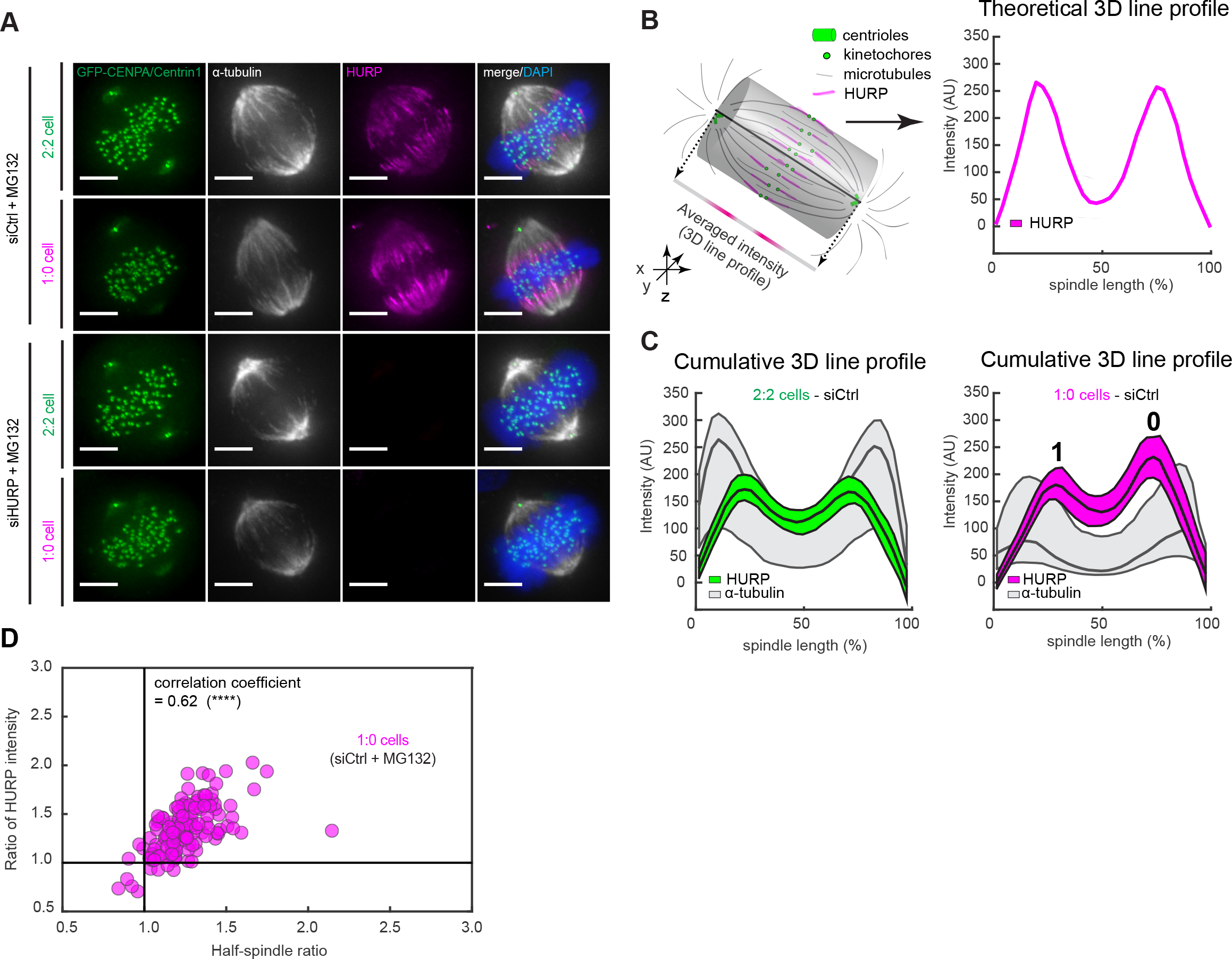
HURP is enriched on the k-fibers of acentrosomal half-spindles in 1:0 cells. (**A**) Immunofluorescence images of control- and HURP-depleted, 2:2 and 1:0 hTert-RPE1 Centrin1-GFP/GFP-CENPA cells blocked in metaphase, stained with α-tubulin and HURP antibodies, and DAPI. Scale bars = 5 μm. (**B**) Schematic representation of how the HURP intensities in the 3D-spindle are transformed into a 1D line HURP-intensity profile along the spindle axis. (**C**) Average HURP intensity line profiles along the spindle axes of all control-depleted 2:2 and 1:0 hTert-RPE1 Centrin1-GFP/GFP-CENPA cells (see Methods section); thick lines – medians; thin lines – 25 and 75 percentiles. N = 3, n = 143-145 cells. (**D**) Spearman correlation graphs of control-depleted 1:0 hTert-RPE1 Centrin1-GFP/GFP-CENPA cells plotting the half-spindle ratios of individual cells and HURP ratios obtained from their spindle 3D line profiles and; N = 3, n = 143-145 cells. **** Indicates p < 0.0001.

The facts that: 1) HURP depletion partially restored spindle symmetry in 1:0 cells (Fig. 3B and C), 2) HURP depletion particularly destabilized k-fibers at acentrosomal half-spindles (Fig. 4E) and 3) HURP was enriched on the k-fibers of the acentrosomal half-spindles (Fig. 5A and C), suggested a possible causal link between spindle asymmetry and HURP asymmetry. Consistent with this hypothesis, we found a positive correlation between the two parameters in 1:0 cells (Spearman correlation coefficient 0.62 respectively; Fig. 5D). The mechanistic origin of HURP asymmetry in 1:0 cells was, nevertheless, unclear.

### HURP localization is linked to spindle asymmetry

To address these mechanisms, we knocked-in by CRISPR/Cas9 an eGFP tag into the *DLGAP5* gene locus encoding for HURP. Sequencing and RNAi experiments confirmed the specificity of eGFP insertion and live cell imaging indicated that the hTert-RPE1 eGFP-HURP cell line showed no alteration in mitotic timing or chromosome segregation efficiency when compared to the parental cell line (Fig. S5A-C; Movie S9). Importantly, 1:0 hTert-RPE1 eGFP-HURP cells also showed an asymmetric HURP distribution (Fig. S5A and D; Movie S10). We first asked whether, once established, HURP asymmetry is static or dynamically maintained during metaphase. To test this, we blocked hTert-RPE1 eGFP-HURP 2:2 or 1:0 cells in metaphase and measured HURP dynamics on the k-fibers by fluorescent recovery after photobleaching (FRAP). We found that the HURP half-life at k-fibers in both 2:2 and 1:0 cells was short (in the order of 10s) and that the initial eGFP-HURP asymmetry was restored after photobleaching (Fig. 6A-F; Fig. S7A-C – note that the apparent large immobile fraction was due to a large fraction of cellular HURP being bleached; Fig. 6G-I). We conclude that HURP asymmetry is actively maintained in metaphase.

**Figure 6.**
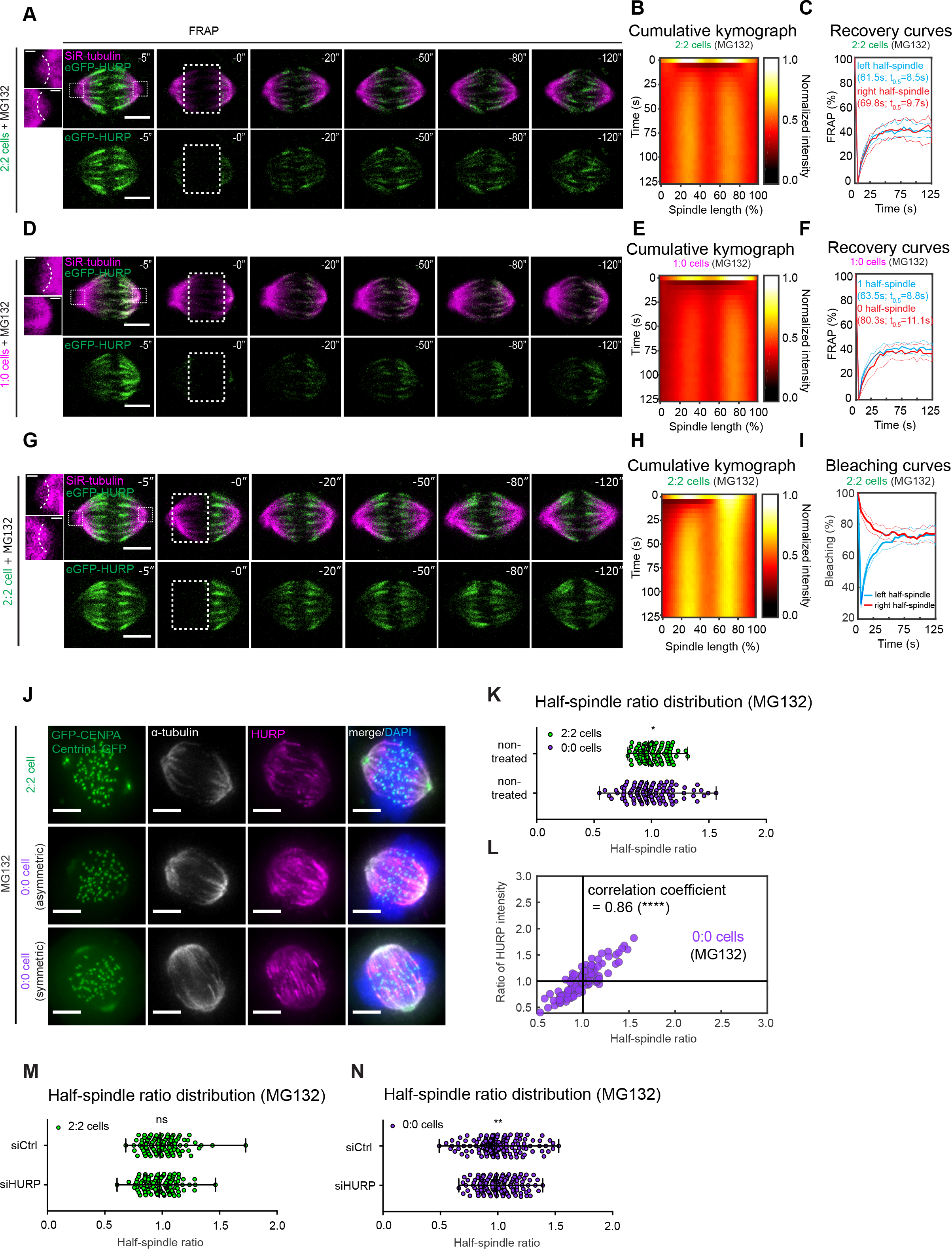
HURP localization is linked to spindle asymmetry. (**A**) Time-lapse images of FRAP performed on both half-spindles of an exemplary 2:2 hTert-RPE1 eGFP-HURP cell stained with SiR-tubulin and blocked in metaphase. Scale bars = 5 μm. Inserts show the morphology of the opposite spindle poles. Scale bars = 1 μm. (**B**) Normalized average local intensity kymograph of HURP line profiles along the spindle axis of all bleached 2:2 cells (see Methods section); N = 3, n = 31 cells. (**C**) Recovery curves of all bleached half-spindles in 2:2 cells; thick lines – means; thin lines – std; N = 3, n = 31 cells. (**D**) Time-lapse images of FRAP performed on both half-spindles of an exemplary 1:0 hTert-RPE1 eGFP-HURP cell stained with SiR-tubulin and blocked in metaphase. Scale bars = 5 μm. Inserts show the morphology of the opposite spindle poles. Scale bars = 1 μm. **(E**) Cumulative kymograph of HURP line profiles along the spindle axis of all bleached 1:0 cells; N = 3, n = 43 cells. (**F**) Recovery curves of all half-spindles bleached in 1:0 cells; thick lines – means; thin lines – s.d.; N = 3, n = 43 cells. (**G**) Time-lapse images of FRAP performed on the left half-spindle of an exemplary 2:2 hTert-RPE1 eGFP-HURP cell stained with SiR-tubulin and blocked in metaphase. Scale bars = 5 μm. Inserts show the morphology of the opposite spindle poles. Scale bars = 1 μm. **(H**) Cumulative kymograph of HURP line profiles along the spindle axis of all bleached 2:2 cells; N = 3, n = 40 cells. (**I**) Bleaching curves of all left and right half-spindles bleached in 2:2 cells; thick lines – means, thin lines – s.d.; N = 3, n = 40 cells. (**J**) Immunofluorescence images of 2:2 and 0:0 hTert-RPE1 Centrin1-GFP/GFP-CENPA cells blocked in metaphase, fixed, stained with α-tubulin and HURP antibodies, and DAPI; Scale bars = 5 μm. (**K**) Half-spindle ratio distributions measured in 2:2 and 0:0 hTert-RPE1 Centrin1-GFP/GFP-CENPA cells; N = 3, n = 108-109 cells; error bars indicate medians with range; Kolmogorov-Smirnov distribution test; p = 0.0236. (**L**) Spearman correlation graphs of 0:0 hTert-RPE1 Centrin1-GFP/GFP-CENPA cells plotting the HURP ratios obtained from the line profiles and half-spindle ratios of individual cells; N = 3, n = 108 cells. (**M**) Half-spindle ratio distributions measured in control- and HURP-depleted 2:2 hTert-RPE1 Centrin1-GFP/GFP-CENPA cells; N = 3, n = 150 cells; error bars indicate medians with range; F-test for variance; p = 0.205. (**N**) Half-spindle ratio distributions measured in control- and HURP-depleted 0:0 hTert-RPE1 Centrin1-GFP/GFP-CENPA cells; N = 3, n = 152-153 cells; error bars indicate medians with range; F-test for variance; p = 0.0023.

We next tested if endogenous HURP localization was under the control of the centrosomal protein kinase Aurora A, which has been proposed to regulate HURP based on HURP overexpression experiments (Wong et al., 2008; Wu et al., 2013; Yu et al., 2005). Immunostaining with antibodies against active Aurora-A (pT288) indicated a higher Aurora-A activity at the centrosomal spindle that could be markedly reduced with 100nM of the Aurora-A inhibitor MLN8237 (Fig. S6A). Nevertheless, Aurora-A inhibition in 2:2 or 1:0 cells did not visibly change HURP localization (Fig. S6B), suggesting that Aurora-A is not the main regulator of HURP localization.

Finally, we prolonged Plk4 inhibition to test in 0:0 cells whether as principle centrosomes regulate HURP localization,. To our surprise, cells without centrosomes displayed a wide range of HURP asymmetries (Fig. 6J and K) that correlated with asymmetric half-spindle ratios (Spearman correlation coefficient 0.86; Fig. 6L). HURP was thereby always enriched on the k-fibers of the shorter half-spindle. This indicated that cells lacking centrosomes fail to balance half-spindles lengths, and that HURP localization is not directly controlled by centrosomes, but rather linked to spindle asymmetry itself. The strong correlation between spindle asymmetry and HURP asymmetry on k-fibers raised the chicken or the egg question, as what come first. HURP-depletion had no effect on spindle (a)symmetry in either 2:2 cells; in contrast in 0:0 cells it did not suppress spindle asymmetry, but significantly reduced the number of outliers (Fig. 6M and N). This suggested that spindle symmetry predates HURP asymmetry, which reinforces spindle asymmetry, as seen in 1:0 cells (Fig. 3B and C).

### HURP localization depends on k-fiber length

In terms of how HURP recognizes spindle asymmetry, we considered two possibilities: differences in k-fiber length and differences in k-fiber dynamics. To distinguish between these hypotheses we used the natural variability of half-spindle lengths in 2:2 hTert-RPE1 Centrin1-GFP/GFP-CENPA cells arrested in metaphase. To specifically test for the contribution of k-fiber dynamics we also treated cells with a low dose of taxol (15 nM), a condition that strongly inhibits k-fiber dynamics (Yvon et al., 1999) but does not disrupt spindle bipolarity (Jordan et al., 1993) (Fig. 7A). We correlated half-spindle length with the length and the overall intensity of the HURP-enrichment zone (“HURP-stripes”) on k-fiber plus-end (see Fig. 7B), and found that shorter half-spindles correlated with shorter but more intense HURP-stripes (Fig. 7C and D). After suppression of k-fiber dynamics with taxol, k-fibers were shorter and bound more HURP; nevertheless HURP intensities were still inverse proportional to half-spindle length (Fig. 7C and D). We conclude that HURP localization depends on k-fiber length, but not k-fiber dynamics. We propose that centrosomes control HURP localization and function at k-fiber plus-end indirectly, by controlling k-fiber length.

**Figure 7.**
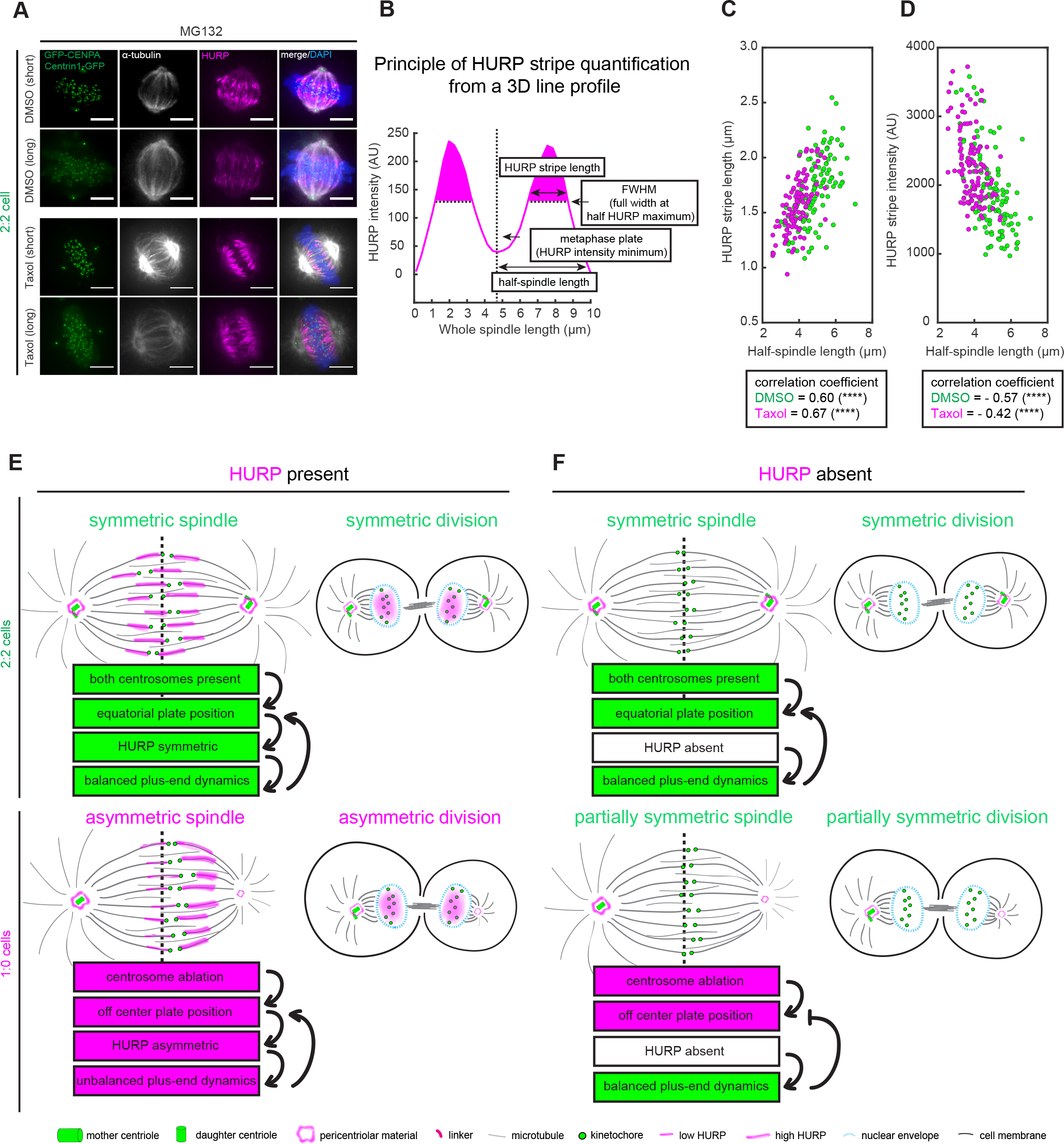
HURP localization depends on k-fiber length. (**A**) Immunofluorescence images of exemplary DMSO-treated (short and long) and taxol-treated 2:2 hTert-RPE1 Centrin1-GFP/GFP-CENPA cells blocked in metaphase, fixed and stained against α-tubulin, HURP and DNA. Scale bars = 5 μm. (**B**) Schematic representation of HURP stripe length and intensity measurement (HURP stripe intensity was calculated as area under the curve limited by the full width at HURP maximum). (**C** and **D**) Spearman correlation graphs of half-spindle length and HURP stripe length (C), or HURP stripe intensity (D) in 2:2 hTert-RPE1 Centrin1-GFP/GFP-CENPA cells treated with DMSO or 15nM of Taxol. **** Indicates p < 0.0001. (**E** and **F**) Speculative model of how centrosomes balance k-fiber plus-end dynamics via HURP, based on our findings in 2:2 (up) and 1:0 (bottom) cells with (E) or without (F) HURP. Briefly, the presence of both centrosomes ensures the equatorial metaphase plate position, which in turn balances HURP and plus-end k-fiber dynamics. Balanced k-fiber plus-end dynamics reinforces spindle symmetry and symmetric divisions (E, top). Asymmetric spindles (mimicked by centrosome ablation) place the metaphase plate off center, which creates k-fibers of unequal lengths and promotes HURP asymmetry. This leads to unbalanced k-fiber plus-end dynamics, which reinforces spindle asymmetry and asymmetric divisions (E, bottom). HURP depletion in cells with both centrosomes does not induce unbalanced k-fiber plus-end dynamics and has no effect on spindle morphology or symmetric divisions (F, top). However, HURP depletion in cells with less than two centrosomes rebalances k-fiber plus-end dynamics, promoting spindle and cell division symmetry (F, bottom).

## DISCUSSION

Centrosomes are dispensable for bipolar spindle assembly in most studied vertebrate cells. However, centrosome loss leads to chromosome segregation errors (Sir et al., 2013), and centrosome manipulation affects k-fiber length (Greenan et al., 2010; Tan et al., 2015), which raises the question as to how centrosomes regulate k-fiber dynamics. Here, we demonstrate that centrosomes balance the dynamics of both k-fiber minus- and plus-ends, ensuring symmetric spindles and divisions. We demonstrate that HURP is one of the mediators of centrosome function, as it regulates plus-end k-fiber dynamics and stability by accumulating on at k-fibers in a manner that is inversely proportional to their length (Fig. 7E and F).

Our data show that single centrosome ablation increases k-fiber stability at the acentrosomal half-spindle, lowers minus-end depolymerization, and increases plus-end pausing, resulting in differential k-fiber behavior between opposite half-spindles. This suggests that, despite the presence of microtubule stabilizing proteins at centrosomes such as MRCS1 and Patronin (Goodwin and Vale, 2010; Hirohashi et al., 2006; Meunier and Vernos, 2011), the main role of the centrosome is to ensure dynamic k-fibers that can rapidly grow and shrink, rather than to stabilize them. Second, we find that bipolar spindles lacking one or both centrosomes are asymmetric. We propose that centrosomes are not mere catalyzers of the bipolar spindle assembly, but that their presence provides balanced k-fiber dynamics at both minus- and plus-ends to ensure spindle symmetry and symmetric divisions. Finally, in agreement with previous studies (Sir et al., 2013) we find that centrosome loss leads to chromosome segregation errors. These errors arose whether k-fiber plus-end dynamics were balanced or not, implying that the transient monopolar configuration is the main cause for formation of erroneous kinetochore-microtubule attachments (Kaseda et al., 2012; Mchedlishvili et al., 2012; Silkworth et al., 2012).

At the molecular level, we show that depletion of microtubule-stabilizing protein HURP (Sillje et al., 2006; Wong and Fang, 2006) restored normal k-fiber plus-end dynamics, and improved spindle and cell division symmetry in 1:0 cells. We therefore propose that centrosomes control k-fiber plus-end dynamics via HURP. This control is however not direct. Instead, we postulate that centrosomes regulate HURP accumulation via k-fiber length, possibly via centrosome-associated proteins such as TPX2 (Bird and Hyman, 2008; Greenan et al., 2010) or TACC family members (Le Bot et al., 2003). Shorter k-fibers accumulate more HURP, suppress plus-end k-fiber dynamics and create a differential k-fiber stability in 1:0 cells (see model in Fig. 7E). Nevertheless, HURP cannot be the only protein by which centrosomes regulated k-fiber dynamics, as its depletion did not normalize poleward microtubule flux or fully rescue spindle symmetry. Finding these missing regulators in the future may be challenging, since the depletion or the inhibition of a number of candidate proteins (e.g. Kif18A, Eg5 or Aurora-A) was not compatible with a stable bipolar spindle in 1:0 cells. Another key future aim will be to determine how HURP “senses” k-fiber length. Indeed, while some kinesins, such as Kif18A selectively accumulate on longer microtubules (Mayr et al., 2007; Varga et al., 2006), this is the first protein that to our knowledge is enriched on shorter microtubules.

Finally, we demonstrate that HURP depletion abolishes the differential k-fiber stability in 1:0 cells, it restores k-fiber plus-end dynamics and leads to more symmetric spindles. This suggests that differential k-fiber dynamics and spindle asymmetry are partially caused by HURP asymmetry. HURP bundles microtubules *in vitro* (Koffa et al., 2006; Santarella et al., 2007; Sillje et al., 2006), which likely leads to k-fiber stabilization *in vivo* (Sillje et al., 2006; Wong et al., 2008). Excessive amounts of HURP on the short acentrosomal k-fibers might over-stabilize their plus-ends in a similar way as microtubule-stabilizing agent taxol “freezes” k-fiber dynamics, resulting in even shorter k-fibers (Snyder and Mullins, 1993; Yvon et al., 1999) (Jordan et al., 1993; Rizk et al., 2009). In addition, by increasing pausing events, HURP accumulation might disrupt the oscillatory sister-kinetochore movements necessary to center the metaphase plate (Tan et al., 2015). We speculate that in normal conditions, apart from its general k-fiber stabilizing activity (Sillje et al., 2006; Wong and Fang, 2006), HURP might specifically stabilize short spindles, to prevent a spindle collapse. The promotion of spindle asymmetry in 1:0 cells would in this case represent an excessive reaction of such a feedback system. Alternatively, in naturally asymmetric spindles (Delaunay et al., 2014; Greenan et al., 2010; Ren and Weisblat, 2006; Roubinet et al., 2017), HURP asymmetry could reinforce the spindle asymmetry by freezing the dynamics of the shorter k-fibers, and thus promote asymmetric cell divisions. This potential function of HURP was not only visible in 1:0 cells, but also in 0:0 cells (Fig. 6J), where the presence of HURP allowed the formation of more extreme spindle asymmetries. HURP is present in humans, mice and frogs (Koffa et al., 2006; Tsou et al., 2003), and related proteins were found in *D. melanogaster* (Zhang et al., 2009) suggesting that the interplay between plus-end k-fiber dynamics and spindle (a)symmetry could be conserved. Altogether, our work identifies a first molecular mechanism by which centrosomes can control spindle function and k-fiber dynamics over large distances.

## METHODS

### Cell culture and drug administration

hTert-RPE1 EB3-GFP/H2B-mCherry (kind gift from W. Krek), hTert-RPE1 Centrin1-GFP/GFP-CENPA (kind gift from A. Khodjakov), and hTert-RPE1 eGFP-HURP cell lines were cultured using DMEM (Thermofisher, Switzerland) medium supplemented with 10 % FCS, 100 U ml^−1^ penicillin and 100 mg ml^−1^ streptomycin (Thermofisher, Switzerland). hTert-RPE1 PA-GFP-α-tubulin cells (Toso et al., 2009) were in addition supplemented with 500 μg ml^−1^ G418 (Life Technologies, Switzerland). All cells were cultured in 37 °C with 5 % CO_2_ in humidified incubators. Live-cell imaging experiments were performed using 8-well ibidi chambers (Vitaris, Switzerland) and imaging medium (Leibovitz L-15 medium from Thermofisher, supplemented with 10 % FCS). The following drug concentrations were used: centrinone 300 nM (kind gift from A. Shiau and Tocris Bioscience), MG132 10 μM (Sigma Aldrich), RO-3306 9 μM (Sigma-Aldrich), SiR-tubulin 25-50 nM (Spirochrome), nocodazole 200 ng ml^−1^ (Sigma Aldrich), SiR-DNA 25 nM (Spirochrome), MLN8237 100nM (Selleckchem), taxol 15 nM (Sigma-Aldrich). All drugs were diluted in the culturing or imaging medium. To obtain 1:0 cells, culturing medium containing freshly thawed centrinone was exchanged twice a day, in order to prevent Plk4 reactivation and centriole over-duplication. All the manipulations with living 1:0 cells (including pretreatment and washing prior fixation) were carried out in the constant presence of centrinone.

### siRNA treatment

Cells were transfected following the provider’s instructions for 32 hours (72 hours in case of Kif2a) with 20 nM siRNAs using Opti-MEM (Thermofisher, Switzerland) and Lipofectamine RNAiMAX (Thermofisher, Switzerland). MEM (Thermofisher, Switzerland) medium supplemented with 10 % FCS was used for transfection. Cells were transferred back to culturing medium 10 hours after siRNA transfection. The following sense strands of validated siRNA duplexes were used: control (Thermofisher, Switzerland) – GGACCUGGAGGUCUGCUGUdTdT (Mchedlishvili et al., 2012); Sas-6 (Thermofisher, Switzerland) – GCACGUUAAUCAGCUACAAdTdT (Leidel et al., 2005); HURP (Qiagen) – GGUGGCAAGUCAAUAAUAAdTdT; HURP alternative sequence (Qiagen) – UGACUCGAUCAGCUACUCAdTdT (Sillje et al., 2006); Kif15 (SMARTpool; GE Healthcare, Switzerland) – GGACAUAAAUUGCAAAUAC, GCAGAGUGUUGAUCAAGAA, GAGCUUCAGUCUUUGCAAA, GAGAUGGAAUGCCUUAGAA (Tanenbaum et al., 2009); Kid-1 (Qiagen) – CAAGCUCACUCGCCUAUUGTT (Wandke et al., 2012); HSET (Qiagen) – UCA GAAGCAGCCCUGUCAA (Cai et al., 2009); CLASP-1 (Qiagen) – GCCAUUAUGCCAACUAUCUdTdT (Mimori-Kiyosue et al., 2005); ch-TOG (Qiagen) – GAGCAGUCGCAAAUGAAGCdTdT (Gergely et al., 2003); Kif2a (ON-TARGETplus, 3796, siRNA-SMARTpools; GE Healthcare, Switzerland) (Tan et al., 2015).

### Live cell imaging

All live-cell imaging experiments were performed using imaging medium at 37°C. To quantify mitotic timing and segregation errors, hTert-RPE1 EB3-GFP/H2B-mCherry cells were imaged for 9-12 h every 3 min with 2 μm z-stacks using either a Nikon Eclipse Ti-E wide-field microscope (Nikon, Switzerland) equipped with a DAPI/eGFP/TRITC/Cy5 filter set (Chroma, USA), a 40X N.A. 1.3 objective, an Orca Flash 4.0 CMOS camera (Hamamatsu, Japan) run with NIS (Nikon). hTert-RPE1 eGFP-HURP and hTert-RPE1 Centrin1-GFP/GFP-CENPA cells were recorded under same conditions using a Olympus DeltaVision wide-field microscope (GE Healthcare, Switzerland) equipped with a eGFP/RFP filter set (Chroma), a 40x 1.3NA/60x 1.4NA objective, and a Coolsnap HQ2 CCD camera (Roper Scientific, USA) run with Softworx (GE Healthcare). To estimate if single centrosome ablation resulted in acute spindle asymmetry hTert-RPE1 EB3-GFP/H2B-mCherry cells were treated with 300 nM centrinone for 32 h, blocked in G2 with 9 μM RO-3306 for 6-7 h, released into a centrinone- and MG132-containing medium, and imaged for 2 h every 5 min. Half-spindle ratio was calculated manually based on 3D stacks using Imaris 7.7 software (BitPlane, Switzerland). Briefly, half-spindle ratio was computed as the length between the center of the mass of the metaphase plate and the EB3 spot at the spindle pole (centrosomal half-spindle) or the end of the diffused EB3 signal around the opposite spindle pole (acentrosomal half-spindle).

### Immunofluorescence

For half-spindle ratio measurements cells were treated with 300 nM centrinone for 32-72 h, 2 h with 10 μM MG132, fixed with 95 % −20 °C methanol for 6 min, rinsed with PBS, blocked with 3 % BSA overnight, and stained with indicated antibodies. To quantify the HURP signal and daughter cells volume cells were fixed in 20 mM PIPES (pH = 6.8), 10 mM EGTA, 1 mM MgCl_2_, 0.2 % Triton X-100, 4 % formaldehyde (Sigma-Aldrich) for 7 min, rinsed with PBS, blocked with 3% BSA overnight, and stained. For HURP quantification at kinetochores cells were rinsed with 5 mM EGTA in PBS, permeabilized using 0.5 % TritonX in PHEM buffer (60 mM Pipes, 25 mM HEPES, 10 mM EGTA, 4 mM MgCl_2_) for 4 minutes, briefly rinsed again with 5 mM EGTA in PBS, fixed with 5 mM EGTA in 95 % −20 °C methanol, for 5 min at RT, and 20 min at −20 °C, then blocked with 3 % BSA and stained. For the cold-stable assay cells were briefly rinsed with ice-cold cytoskeleton buffer (CB buffer; 10 mM MES, 150 mM NaCl_2_, 5 mM MgCl_2_, 5 mM glucose; Sigma-Aldrich), fixed with an ice-cold glutaraldehyde-based solution in CB buffer (0.05 % glutaraldehyde, 3 % formaldehyde and 0.1 % TritonX; Sigma Aldrich) for 15 min on ice, rinsed twice with CB buffer for 10 min at RT, rinsed once with PBS, blocked with 3 % BSA for 30 min and stained. The following primary antibodies were used: rabbit serum anti-γ-tubulin (1:2000; this study), mouse monoclonal anti-γ-tubulin (1:1000; Sigma-Aldrich T6557; clone GTU-88), rabbit serum anti-HURP (1:500; kind gift from E. Nigg; (Sillje et al., 2006)), monoclonal mouse anti-α-tubulin (1:500; Sigma-Aldrich; T9026; clone DM1A), rabbit polyclonal anti-α-tubulin (1:1000; abcam; ab1851), mouse monoclonal anti-pericentrin (1:1000; abcam; ab28144), rabbit monoclonal anti-Aurora-A-T288 (1:500; Cell Signaling; C39D8 clone). To stain cell membranes, cells were incubated for 15 min with Cell Mask (1:1000; Thermofisher Switzerland) in PBS before mounting the coverslips with VECTASHIELD with DAPI (Vector Laboratories). Cross-absorbed secondary anti-mouse and anti-rabbit antibodies (Invitrogen) were used. For half-spindle ratio and HURP localization measurements, z-stack images were taken using an Olympus DeltaVision wide-field microscope (GE Healthcare, Switzerland) equipped with a eGFP/RFP filter set (Chroma) and with 60x NA 1.4 or 100X 1.4 NA objectives and recorded with a Coolsnap HQ2 CCD camera (Roper Scientific, USA) and the Softworx software (GE Healthcare) with 0.2 μm spacing between stacks. For the quantification of HURP intensity at kinetochore microtubule plus-ends, the images were taken using a laser scanning LSM800 confocal microscope (Zeiss) with an Airyscan function, 63X NA 1.4 Oil DIC f/ELYRA objective with 405/488/561/640nm lasers and Airyscan-mode optimized settings (final pixel size of 35 nm in x and y, and 190 nm in z axis). Images of 3 μm thickness were 3D-deconvolved using ZEN software (Zeiss) and analyzed in MATLAB 2017b (The MathWorks, Inc. Natick MA, USA).

### Image processing and analysis

To calculate the half-spindle ratio, 3D images were deconvolved using Softworx (GE Healthcare) and reconstructed using Imaris 7.7 (BitPlane, Switzerland). The length of the opposite half spindles was defined as the 3D distance between the spindle poles (based on γ-tubulin staining or centrin1-GFP signal) and the center of the metaphase plate (computed using the “surface” function and DAPI staining). For 2:2 cells, the spindle symmetry was calculated as the ratio between the lengths of the half-spindle containing the brightest centriole and the length of the opposite half-spindle. For 1:0 cells, the spindle poles were assigned based on the y-tubulin staining, and the spindle asymmetry was defined as the ratio between the lengths of the centriole-bearing half-spindle and the opposite half-spindle. To quantify the symmetry of cell division, the “surface” function in Imaris 7.7 software (BitPlane, Switzerland) was used. Cell Mask staining enabled defining cell borders and then automatic volume computation. The ratio was calculated dividing the volume of the daughter cell containing the brightest centriole (2:2 cells) or the only centriole (1:0 cells) by the volume of the opposite daughter cell.

### Kinetochore microtubule depolymerization assay

hTert-RPE1 Centrin1-GFP/GFP-CENPA cells were treated in parallel with control or HURP-siRNA and with DMSO or 300 nM centrinone for 32 h. Prior to imaging, 25 nM SiR-tubulin was added for 4 h and 10 μM MG132 for 1 h. Metaphase 1:0 cells were selected based on the presence of a single centriole-bearing spindle pole in the SiR-tubulin channel. Medium containing 200 ng ml^−1^ nocodazole, 300 nM centrinone, 25 nM SiR-tubulin and 10 μM MG132 was added on the microscope stage upon launching live cell imaging. Series of 3D z-stacks were acquired with 0.5 μm spacing every 1 min for 15 min using an Olympus DeltaVision wide-field microscope (GE Healthcare, Switzerland) equipped with a eGFP/RFP filter set (Chroma) a 100x NA 1.4 objective, a Coolsnap HQ2 CCD camera (Roper Scientific, USA) run by Softworx (GE Healthcare). The integrities of the opposite half-spindles were monitored from the moment from nocodazole addition until a given half-spindle collapsed, and lost its connection to the spindle pole (which roughly corresponds to the class 3 in the cold-stable assay, see main text). For whole spindle intensity decay quantification, 4D images (xyzt) of each cell were segmented relying on Otsu’s method and maximum intensity projection along the z-axis for all the time points using a custom-made framework developed with MATLAB 2016a (MathWorks Natick MA USA). Thereafter, the value of the pixels belonging to the cell of interest was summed, resulting in a fluorescence intensity signal over time. Microtubule loss was expressed as a percentage of the decay of the SiR-tubulin signal normalized to the first time point (100%). Statistical analysis of microtubule depolymerization was done using repeated measures analysis of variance with Greenhouse-Geisser correction of the sphericity followed by a post-hoc multiple comparison by time.

### Kinetochore tracking assay

hTert-RPE1 Centrin1-GFP/GFP-CENPA cells were treated for 32h in parallel with either control or HURP siRNA in combination with DMSO or 300 nM centrinone. Metaphase cells were imaged for 5 min every 7.5 s using an Olympus DeltaVision wide-field microscope (GE Healthcare, Switzerland) equipped with a eGFP/RFP filter set (Chroma), a 60x NA 1.4 objective and a Coolsnap HQ2 CCD camera (Roper Scientific, USA) run under Softworx (GE Healthcare) with 0.5 μm z-stacks. The images were cropped and deconvolved in Softworx, and analyzed using an automated KT tracking code written in MATLAB 2013b (The Math Works, Inc, Natic, USA). The latest code is available under https://github.com/cmcb-warwick. The output of this analysis is the frame-to-frame displacement of sister-KTs and their relative distance from the center of the metaphase plate. We used an auto-correlation function to quantify the regularity of the sister-kinetochore oscillations along the spindle axis. To measure the sister-kinetochore pausing frequency and the mean sister-KT pair displacement, we fitted normal probability density function to the data using maximum likelihood estimation (Olziersky et al., 2018).

### Poleward microtubule flux measurement

hTert-RPE1 PA-GFP-α-tubulin cells were incubated with 300 nM centrinone for 32 h, 50 nM SiR-tubulin in the last 2 h and 10 μM MG132 in the last 30 min. The cells incubated in MG132 were imaged for no longer than 2 h. Metaphase 1:0 cells were selected based on the presence of a single centriole-bearing spindle pole in the SiR-tubulin channel. Single focal planes of 150 nm pixel size were acquired using an A1r point scanning confocal microscope (Nikon), 60X 1.4 NA CFI Plan Apochromat objective and NIS elements software. Both half-spindles were photo-activated at the same time with a 500 ms 100 % 405 nm laser pulse using a 1 pixel-thick and 11 um-long ROI stretched across the spindle. Single focal planes were imaged every 5 s for 1 min. Photo-activated kinetochore-microtubule bundles were detected using an in-build “spot” function (500 μm diameter) and Imaris 7.7 (Bitplane). Spots were tracked for 60 s and the distance between each spot and each spindle pole was computed for each frame. To ensure accurate measurements, a single k-fiber was tracked per half-spindle. Final flux rates per half-spindle were calculated as the distance, at which the spot traveled towards its spindle pole within 1 min.

### HURP line profiling and FRAP experiments

Imaris 8.2 (Bitplane, Switzerland) with custom-made extensions written in MATLAB 2017b (MathWorks, Natick MA USA) were used to transform the 3D HURP signal into a 1D line profile along the spindle axis. Briefly, spindle poles were automatically or manually detected using the centrin signal, and the images were isotopically rescaled proportionally to spindle length. This provided an axis connecting the opposite poles and allowed building either a 3D right cylinder (see Figure 5X). Thereafter the generated volume was cut in 33 slices along the spindle axes and the average fluorescence per voxel was extracted in each slice to build-up an intensity line-profile along the spindle. The profile orientation was decided either manually or randomly assigned. For 3D line profiling, signals coming from HURP and α-tubulin channels were background subtracted using the 10^th^ percentile value within the cylinder. The metaphase plate position was inferred from the maximum DAPI value position within the cylinder; this allowed finding the mean intensity of HURP and α-tubulin for each half-spindle. We proceed the same way for FRAP experiment except that in this case we started from a 2D surface instead of a 3D cylinder, and the poles were detected based on the SiR-tubulin signal. HURP ratio was computed as the ratio of the two half-spindles intensities (more intense / less intense). Depletion efficiency per half-spindle was computed by dividing the average HURP intensity of a given half-spindle (left or right for 2:2 cells; 1 or 0 for 1:0 cells) of HURP-depleted cells by the average intensity of HURP in a respective half-spindle from control-depleted cells. For correlation of HURP intensity ratio and half-spindle ratio, the Spearman correlation value was computed using the automatic HURP intensity ratio provided by the 3D line profiling tool and the half-spindle ratio calculated manually using Imaris 8.2 (BitPlane, Switzerland; see above).

### Generation of the hTert-RPE1 eGFP-HURP knock-in cell line

A pCMV-Cas9-RFP vector encoding both the Cas9 gene and a guide RNA (gRNA) was designed and provided by Sigma-Aldrich. The gRNA targeted the ACATTTTGCCAGTCGACACAGG sequence within the first exon of the human *DLGAP5* gene encoding the HURP protein. The PAM sequence was located 33 base pairs downstream of the start codon. The repair template was designed using SnapGene software (GSL Biotech) such that it contained 700 bp overhangs flanking the Cas9-cutting site (ACATTTTGCCAGTCGA<u>CA</u>CAGG), which included the genomic DNA, a spliced sequence encoding for full length eGFP and a linker of five glycins between eGFP and HURP. The PAM sequence in the repair template was modified to prevent Cas9 from cutting the construct after homologous recombination. The repair template was synthesized and cloned into the pUC57-Kan vector by GENEWIZ. hTert-RPE1 cells were co-transfected for 24 h with pCMV-Cas9-RFP and pUC57-kan plasmids using Opti-MEM and Lipofectamine 2000 (Thermofisher Switzerland) following the provider’s instructions. Cells were kept in culture for 6 days, sorted for GFP-positive cells (0.0007 % of the entire population), and cultured for another 6 days as a mixed population in conditioned medium (mix of 50 % fresh culturing medium and 50 % filtered medium harvested from a confluent culture of hTert-RPE1 parental cell line). Single GFP-positive and RFP-negative cells (about 50 % of the entire population; no RFP-positive cells were observed) were sorted into a 96-well plate and cultured in conditioned medium. 96-well plate was screened for eGFP clones with a localization resembling HURP localization using a Nikon Eclipse Ti-E wide-field microscope (Nikon, Switzerland) equipped with a DAPI/eGFP/TRITC/Cy5 filter set (Chroma, USA) and an air 40X NA 1.3 objective. Positive clones were tested by genomic DNA sequencing (Supplementary materials) and their mitotic behavior assessed. Clone F8 was chosen based on a correct eGFP insertion, normal mitotic timing and chromosome segregation rates comparable to the parental cell line. siHURP treatment resulted in an efficient depletion of the eGFP protein in this clone confirming the presence of eGFP-HURP fusion protein.

### Fluorescence recovery after photobleaching

hTert-RPE1 eGFP-HURP cells were treated either with DMSO or 300 nM centrinone for 32 h and incubated prior imaging with 25 nM SiR-tubulin for 4 h and 10 μM MG132 for 30 min. Metaphase 1:0 monoastral cells were selected based on the SiR-tubulin signal and strong HURP asymmetry. Single focal planes of 120 nm pixel size were acquired using an A1r point scanning confocal microscope (Nikon), 60X 1.4 NA CFI Plan Apochromat objective and NIS elements software. An ROI of 7 μm x 11 μm enclosing an area around the metaphase plate or one of the half-spindle spindles was bleached using a 500 ms pulse of 100% 405 nm laser. The bleached cells were imaged every 2 s for 2 min. The acquired images were processed using a modified 3D line-profiling tool (see above). Briefly, spindle poles were automatically detected based on the SiR-tubulin signal, and tracked over time to provide an axis connecting the opposite poles to correct for spindle rotation. The generated 2D surface of 8 μm width along the long spindle axis was then sliced in 33 steps along the pole-to-pole axis. Fluorescent average pixel values were computed in each slice creating an intensity line-profile along the spindle axis. The profile orientation was decided either manually (1:0 cells) or randomly assigned (2:2 cells). The intensity profiles were corrected for photobleaching, using intensity profiles of unbleached spindles. In each half-spindle, the mean maximal fluorescence intensity loss in the HURP channel was fitted with a double exponential decay: 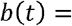 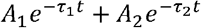. The FRAP efficiency, corrected for the photobleaching due to imaging, was expressed as 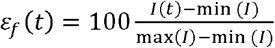, where I is the maximum fluorescent intensity in a given half-spindle, i.e. indicating the localization of HURP, and min(I) and max(I) are respectively the values just before and just after the bleaching. The bleaching efficiency was computed as: 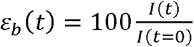.

## SUMMARY OF SUPPLEMENTARY MATERIAL

The supplementary materials contain six supplementary figures supporting the main figures and ten supplementary movies.

## Acknowledgements

The authors thank A. Khodjakov (NY State University, USA) and A. Shiau (Ludwig Institute for Cancer Research, CA, USA) for reagents; A. Eskat for producing the anti-γ-tubulin antibody; the Bioimaging Facility at the Medical Faculty of the University of Geneva for help in data analysis; A. McAinsh (University of Warwick, UK), S. Royle (University of Warwick, UK), M. Gotta, A-M. Olziersky, T. Wilhelm and C. Castrogiovanni (all University of Geneva, Switzerland) for critical discussions of the manuscript. Work in the Meraldi laboratory is supported by the SNF-project grants (No 31003A_160006 and 31003A-179413) and the University of Geneva. The authors declare no competing financial interests.

## Authors contributions

The project was initiated by Damian Dudka and Patrick Meraldi and directed by P.M. D.D. performed all the experiments. Nicolas Liaudet wrote the codes to measure HURP localization. Hélène Vassal wrote the code to measure whole spindle intensity decay. D.D. and P.M. analyzed and interpreted all the results with contribution from N.L. and H.V. D.D and P.M. wrote the manuscript.

## Data availability

All data are available upon request. Due to the large size of the raw images and movies, these data will require transfer to a hard drive.

## Code availability

Codes measuring HURP localization and the whole spindle intensity decay are available upon request.

## Movie S1

Time-lapse movie of control-depleted hTert-RPE1 EB3-eGFP/H2B-mCherry cell treated for 32 h with DMSO, showing mitotic progression. Images were taken every 3 min for 12 hours using a wide-field microscope. Scale bars = 5 μm; green – EB3-eGFP; magenta – H2B-mCherry (see Fig. 1B).

## Movie S2

Time-lapse movie of control-depleted hTert-RPE1 EB3-eGFP/H2B-mCherry cell treated for 32 h with 300 nM of centrinone, showing mitotic progression. Images were taken every 3 min for 12 hours using a wide-field microscope. Scale bars = 5 μm; green – EB3-eGFP; magenta – H2B-mCherry (see Fig. 1B).

## Movie S3

Time-lapse movie of control-depleted 2:2 hTert-RPE1 Centrin1-GFP/GFP-CENPA metaphase cell, showing sister-KT oscillations. Images were taken every 7.5 s for 5 min using a wide-field microscope; green – GFP-CENPA (see Fig. 2G).

## Movie S4

Time-lapse movie of control-depleted 1:0 hTert-RPE1 Centrin1-GFP/GFP-CENPA metaphase cell, showing sister-KT oscillations. Images were taken every 7.5 s for 5 min using a wide-field microscope; green – GFP-CENPA (see Fig. 2G).

## Movie S5

Time-lapse movie of HURP-depleted 2:2 hTert-RPE1 EB3-eGFP/H2B-mCherry cell, showing mitotic progression. Images were taken every 3 min for 12 hours using a wide-field microscope. Scale bars = 5 μm; green – EB3-eGFP; magenta – H2B-mCherry (see Fig. S2E).

## Movie S6

Time-lapse movie of HURP-depleted 1:0 hTert-RPE1 EB3/H2B-mCherry cell, showing mitotic progression. Images were taken every 3 min for 12 hours using a wide-field microscope. Scale bars = 5 μm; green – EB3-eGFP; magenta – H2B-mCherry (see Fig. S2E).

## Movie S7

Time-lapse movie of HURP-depleted 2:2 hTert-RPE1 Centrin1-GFP/GFP-CENPA metaphase cell, showing sister-KT oscillations. Images were taken every 7.5 s for 5 min using a wide-field microscope; green – GFP-CENPA (see Fig. 4H).

## Movie S8

Time-lapse movie of HURP-depleted 1:0 hTert-RPE1 Centrin1-GFP/GFP-CENPA metaphase cell, showing sister-KT oscillations. Images were taken every 7.5 s for 5 min using a wide-field microscope; green – GFP-CENPA (see Fig. 4H).

## MovieS9

Time-lapse movie of a 2:2 hTert-RPE1 eGFP-HURP cell incubated with 25 nM of SiR-tubulin, showing mitotic progression. Images were taken every 3 min for 9 hours using a wide-field microscope. Scale bars = 5 μm; green – eGFP-HURP; magenta – SiR-tubulin (see Fig. S5A).

## Movie S10

Time-lapse movie of a 1:0 hTert-RPE1 eGFP-HURP cell incubated with 25 nM of SiR-tubulin, showing mitotic progression. Images were taken every 3 min for 9 hours using a wide-field microscope. Scale bars = 5 μm; green – eGFP-HURP; magenta – SiR-tubulin (see Fig. S5A).

