## Supplementary Files for "Centrosomes control kinetochore-fiber plus-end dynamics via HURP to ensure symmetric divisions"

### Dudka et al., Figure S1 (Related to Figure 1).

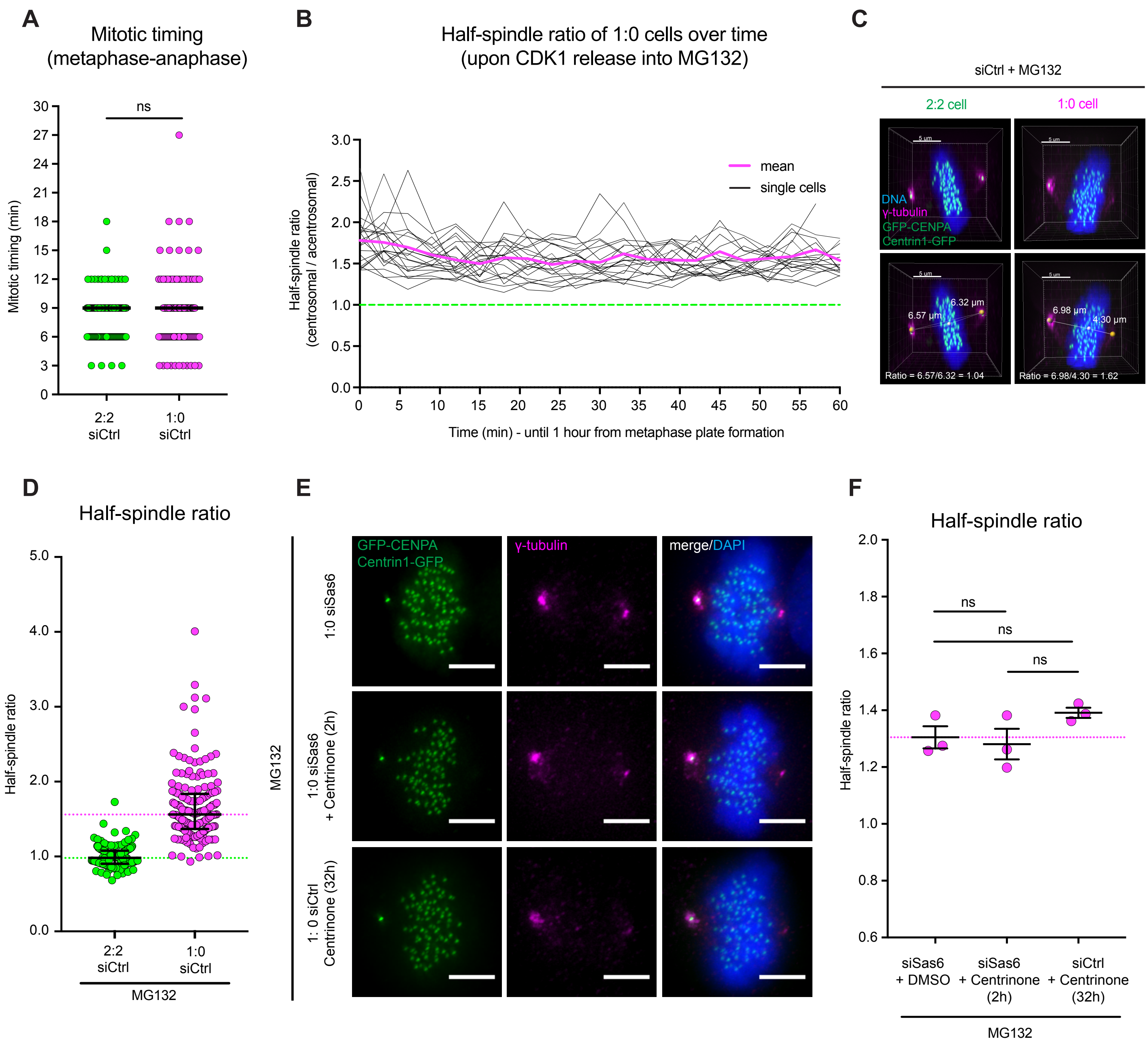

**Figure S1 (Related to Figure 1). Centrosome ablation leads to asymmetric spindles and asymmetric divisions** (**A**) Time from metaphase plate formation to anaphase onset measured in control-depleted hTert-RPE1 EB3-GFP/H2B-mCherry cells treated for 32 h with either DMSO (green) or 300 nM centrinone (magenta); N = 4; n =135-138 cells; error bars show median and 95% CI; Mann-Whitney test; p = 0.5162. (**B**) Half-spindle ratio measured over time in hTert-RPE1 EB3-eGFP/H2B-mCherry 1:0 cells released from a Cdk1-inhibtion block, and re-blocked in metaphase with MG132. Green dashed line represents ratio = 1 (perfect symmetry); N = 4, n = 22 cells. (**C**) 3D reconstitution of immunofluorescence images of control depleted 2:2 and 1:0 hTert-RPE1 Centrin1-GFP/GFP-CENPA metaphase cells, stained with γ-tubulin antibodies and DAPI. Scale bars = 5 μm. The images show how half-spindle ratios were quantified in 3D. The spots (orange = spindle poles; white = metaphase plate center of the mass) were assigned automatically. (**D**) Spread of half-spindle ratios in single control-depleted 2:2 and 1:0 hTert-RPE1 Centrin1-GFP/GFP-CENPA cells; error bars represent median and 95% CI; N = 3; n = 150-169 cells; statistics reported in Fig. 1E. (**E**) Immunofluorescence images of hTert-RPE1 Centrin1-GFP/GFP-CENPA cells transfected with control- or Sas6- siRNA for 32 h and co-incubated with either DMSO or 300 nM centrinone for 2 or 32 h, and blocked in metaphase prior fixation. Scale bars = 5 μm. (**F**) Quantification of the half-spindle ratio in control- or Sas6-depleted 1:0 hTert-RPE1 Centrin1-GFP/GFP-CENPA cells treated with centrinone for indicated times and blocked in metaphase prior fixation; dashed line shows the mean half-spindle ratio for Sas6-depleted and DMSO-treated cells; N = 3, n = 112-141 cells; error bars represent mean and s.e.m.; siSas6 + DMSO vs. siSas6 + centrinone (2h) p = 0.8311 ; siSas6 + DMSO vs. siCtrl + centrinone (32h) p = 1979 ; siSas6 + centrinone (2h) vs. siCtrl + centrinone (32h) p = 0.1085 ; in two-tailed unpaired t-test; ns – not significant. * Indicates p < 0.05, ns – not significant.

**Dudka et al., Figure S2 (Related to Figure 3).**

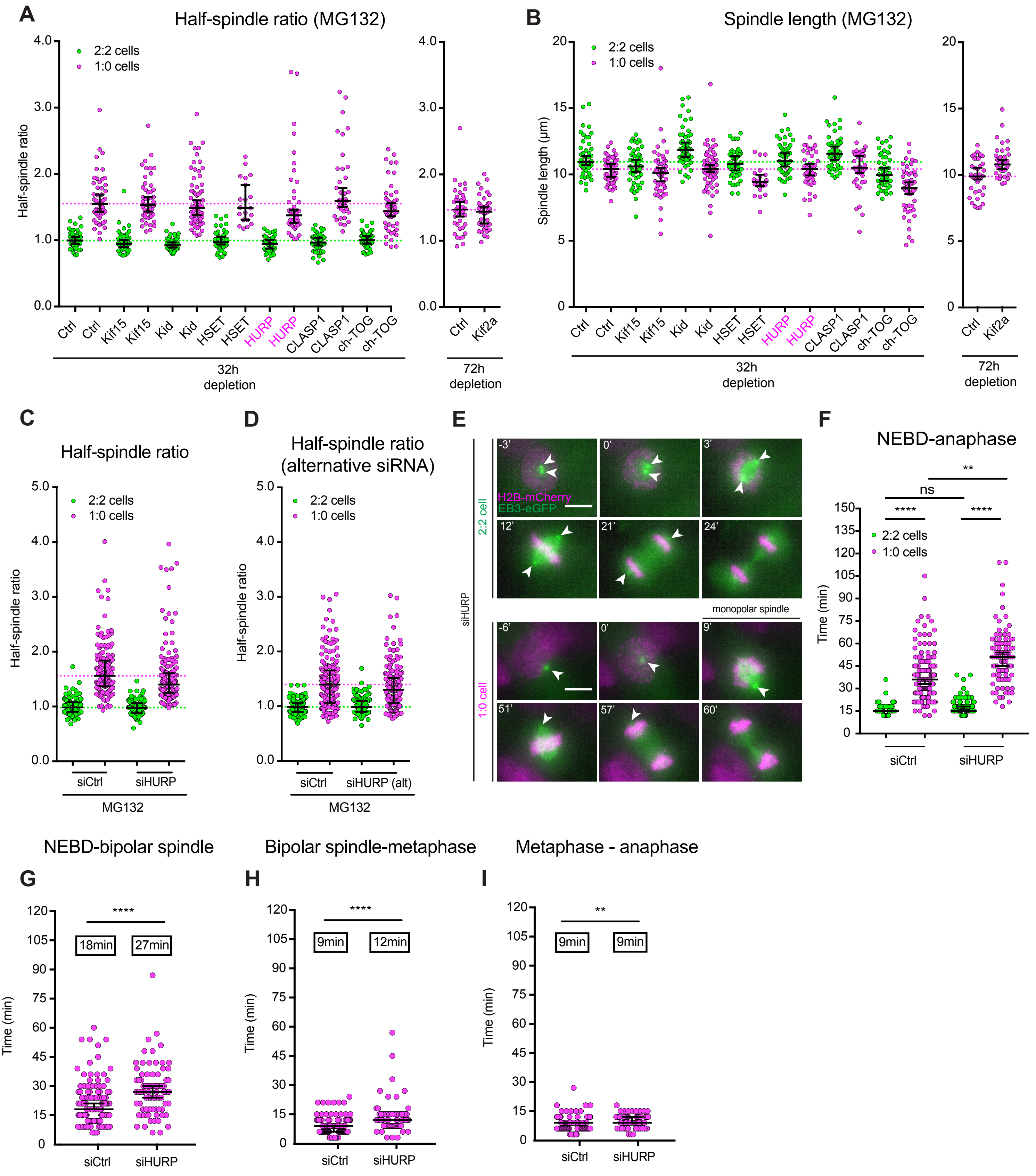

**Figure S2 (Related to Figure 3). HURP depletion partially rescues asymmetric spindles and asymmetric divisions A**) Half-spindle ratios of single 2:2 and 1:0 hTert-RPE1 Centrin1-GFP/GFP-CENPA cells treated with non-targeting or indicated targeting siRNA; n = 20-72 cells. Dashed lines show median half-spindle ratio for the control-depleted 2:2 cells (green) and 1:0 cells (magenta). Efficient Kif2a depletion required 72h treatment (Tan et al., 2015). (**B**) Spindle length of single 2:2 and 1:0 hTert-RPE1 Centrin1-GFP/GFP-CENPA cells transfected with the non-targeting or indicated targeting siRNAs, blocked in metaphase prior fixation. Dashed lines represent the median spindle length for 2:2 (green) or 1:0 (magenta) cells treated with the non-targeting siRNA; n = 20-72 cells. (**C**) Spread of half-spindle ratios in single control- or HURP-depleted, 2:2 and 1:0 hTert-RPE1 Centrin1-GFP/GFP-CENPA cells; error bars represent median and 95% CI; N = 3; n = 150-176 cells; statistics reported in Fig. 3B. (**D**) Spread of half-spindle ratios after control depletion or depletion with an alternative siRNA oligonucleotide set; error bars represent median and 95% CI; N = 4; n = 180-181 cells; statistics reported in 3C. (**E**) Time-lapse images of HURP-depleted 2:2 and 1:0 hTert-RPE1 EB3/H2B-mCherry cells. White arrowheads point at centrosomes. Scale bars = 5 µm; (see Supplementary Movies 5 and 6). (**F**) Mitotic timing measured from NEBD until anaphase onset in control- and HURP-depleted, 2:2 and 1:0 hTert-RPE1 EB3/H2B-mCherry cells; N = 4, n = 95-182 cells; error bars represent median and 95% CI; 2:2 siCtrl vs. 1:0 siCtrl $p < 0.0001$; 2:2 siHURP vs. 1:0 siHURP $p < 0.0001$; 2:2 siCtrl vs. 2:2 siHURP $p = 0.4571$; 1:0 siCtrl vs. 1:0 siHURP $p = 0.001$ in Kruskal-Wallis test. (**G**) Time from NEBD to bipolar spindle formation in 1:0 cells; N = 4, n = 76-136 cells; error bars represent median and 95% CI; Mann-Whitney test; $p < 0.0001$. (**H**) Time from bipolar spindle formation to metaphase in 1:0 cells; N = 4, n = 75-134 cells; error bars represent median and 95% CI; Mann-Whitney test; $p < 0.0001$. (**I**) Time from metaphase to anaphase onset in 1:0 cells; N = 4, n = 75-135 cells; error bars represent median and 95% CI; Mann-Whitney test; $p = 0.0046$. ** Indicates $p < 0.01$; **** indicates $p < 0.0001$; ns – not significant.

### Dudka et al., Figure S3 (Related to Figure 4).

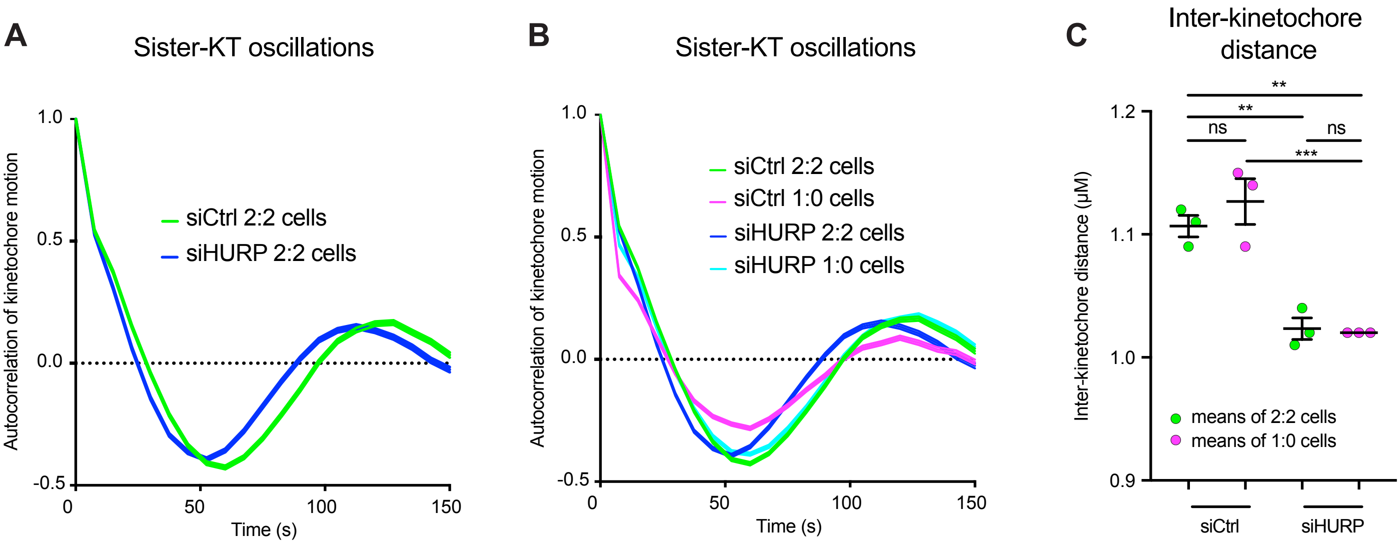

**Figure S3 (Related to Figure 4). HURP depletion rescues plus-end k-fiber dynamics** (**A**) Cumulative autocorrelation curves representing sister-KT pair oscillations in 2:2 control-depleted (green curve) and 2:2 HURP-depleted (blue curve) hTert-RPE1 Centrin1-GFP/GFP-CENPA cells; N = 3, n = 36 cells; $n_{kt}$ = 1289-1457 pairs. Line thickness indicates s.d. (**B**) Merge of the cumulative autocorrelation curves from of control- and HURP-depleted 2:2 and 1:0 hTert-RPE1 Centrin1-GFP/GFP-CENPA cells. (**C**) Quantification of the inter-kinetochore distances from control- or HURP-depleted 2:2 and 1:0 hTert-RPE1 Centrin1-GFP/GFP-CENPA cells. N = 3, n = 36 cells; $n_{kt}$ = 1289-1457 pairs. ** Indicates $p < 0.01$; *** indicates $p < 0.001$; ns – not significant.

### Dudka et al., Figure S4 (Related to Figure 5)

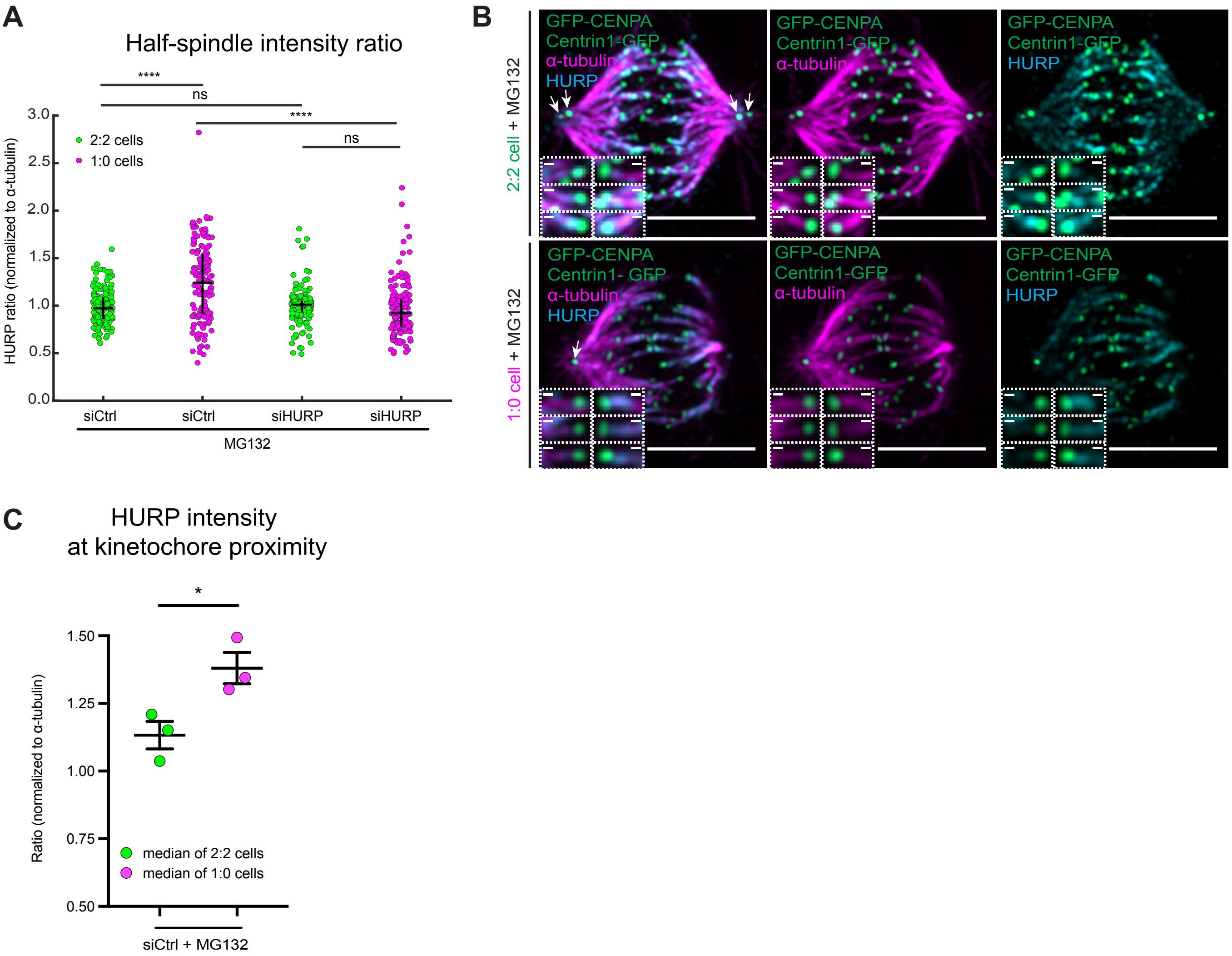

**Figure S4 (Related to Figure 5). HURP asymmetry is not due to tubulin asymmetry** (**A**) Half-spindle HURP intensity ratios of single control- and HURP-depleted, 2:2 and 1:0 hTert-RPE1 Centrin1-GFP/GFP-CENPA cells quantified based on single HURP line profiles along the spindle axis; N = 3, n = 145-151 cells; 2:2 siCtrl vs. 1:0 siCtrl $p < 0.0001$; 2:2 siHURP vs. 1:0 siHURP $p = 0.6189$; 2:2 siCtrl vs. 2:2 siHURP $p = 9656$; 1:0 siCtrl vs. 1:0 siHURP $p < 0.0001$ in two-way ANOVA. (**B**) Immunofluorescence images of control-depleted 2:2 and 1:0 hTert-RPE1 Centrin1-GFP/GFP-CENPA cells blocked in metaphase prior fixation, stained with α-tubulin and HURP antibodies. Scale bars = 5 μm. White arrows indicate centrioles. Inserts show examples of HURP localization at the proximity of single kinetochore microtubule bundles. Scale bars = 0.25 μm. (**D**) Quantification of the ratio of kinetochore-proximal HURP intensity between k-fibers from opposite half-spindles, normalized to α-tubulin signal; N = 3; n = 30 cells; $n_{kt\text{-}bundles} = 1077\text{-}1098$; error bars show mean and s.e.m.; two-tailed unpaired t-test; $p = 0.0323$. * Indicates $p < 0.05$, **** indicates $p < 0.0001$; ns – not significant.

### Dudka et al., Figure S5 (Related to Figure 6)

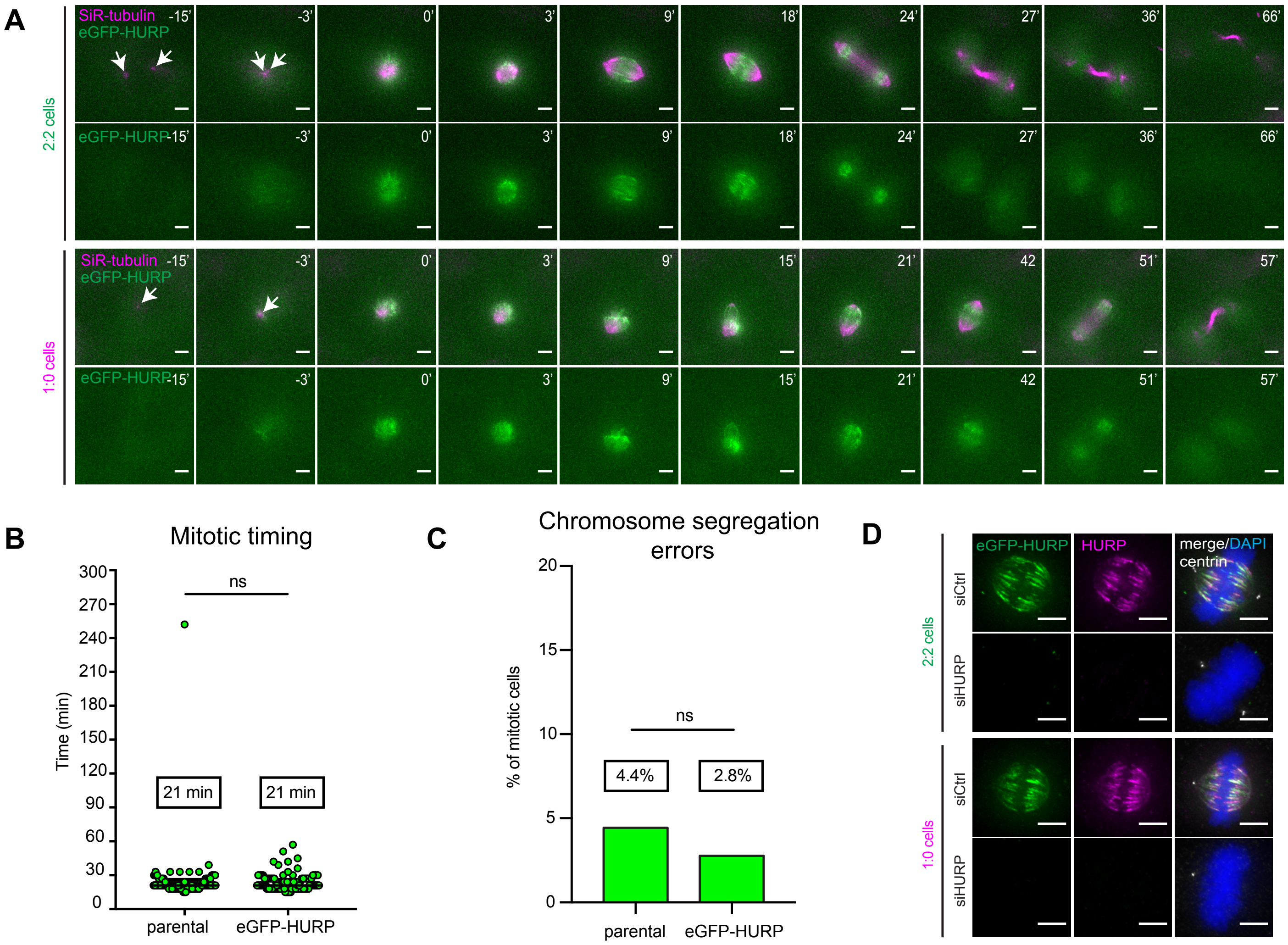

**Figure S5 (Related to Figure 6)**. **eGFP-HURP is asymmetric in living 1:0 cells** (**A**) Time-lapse images of 2:2 and 1:0 hTert-RPE1 eGFP-HURP cells incubated with 25nM SiR-tubulin and recorded for 9 hours in 3 min intervals. Arrows indicate centrosomal spindle poles. Scale bars = 5 µm; (see Supplementary Movies 9 and 10). (**B**) Mitotic timing measurement of non-treated hTert-RPE1 (parental) and hTert-RPE1 eGFP-HURP cells incubated with 25 nM SiR-DNA and imaged for 12h with in 3 min intervals; N = 3, n = 135-144 cells; Mann-Whitney test; p = 0.5479. **(C**) Quantification of segregation errors in non-treated hTert-RPE1 (parental) and hTert-RPE1 eGFP-HURP cells; N = 3, n = 136-144 cells; Fisher's exact test with 99% CI; p = 0.5303. (**D**) Immunofluorescence images of control- or HURP-depleted 2:2 and 1:0 hTert-RPE1 eGFP-HURP cells stained for HURP and DNA. Scale bars = 5 µm. ns – not significant.

### Dudka et al., Figure S6 (Related to Figure 6)

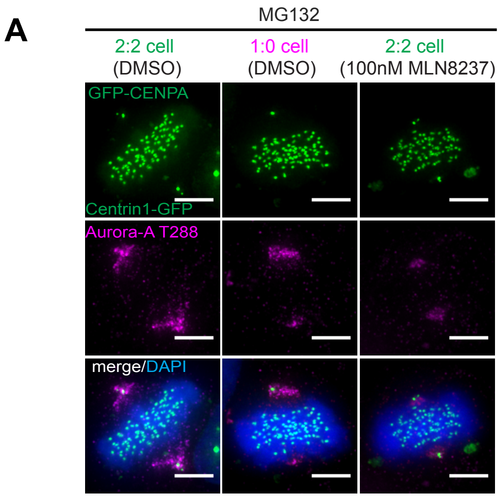


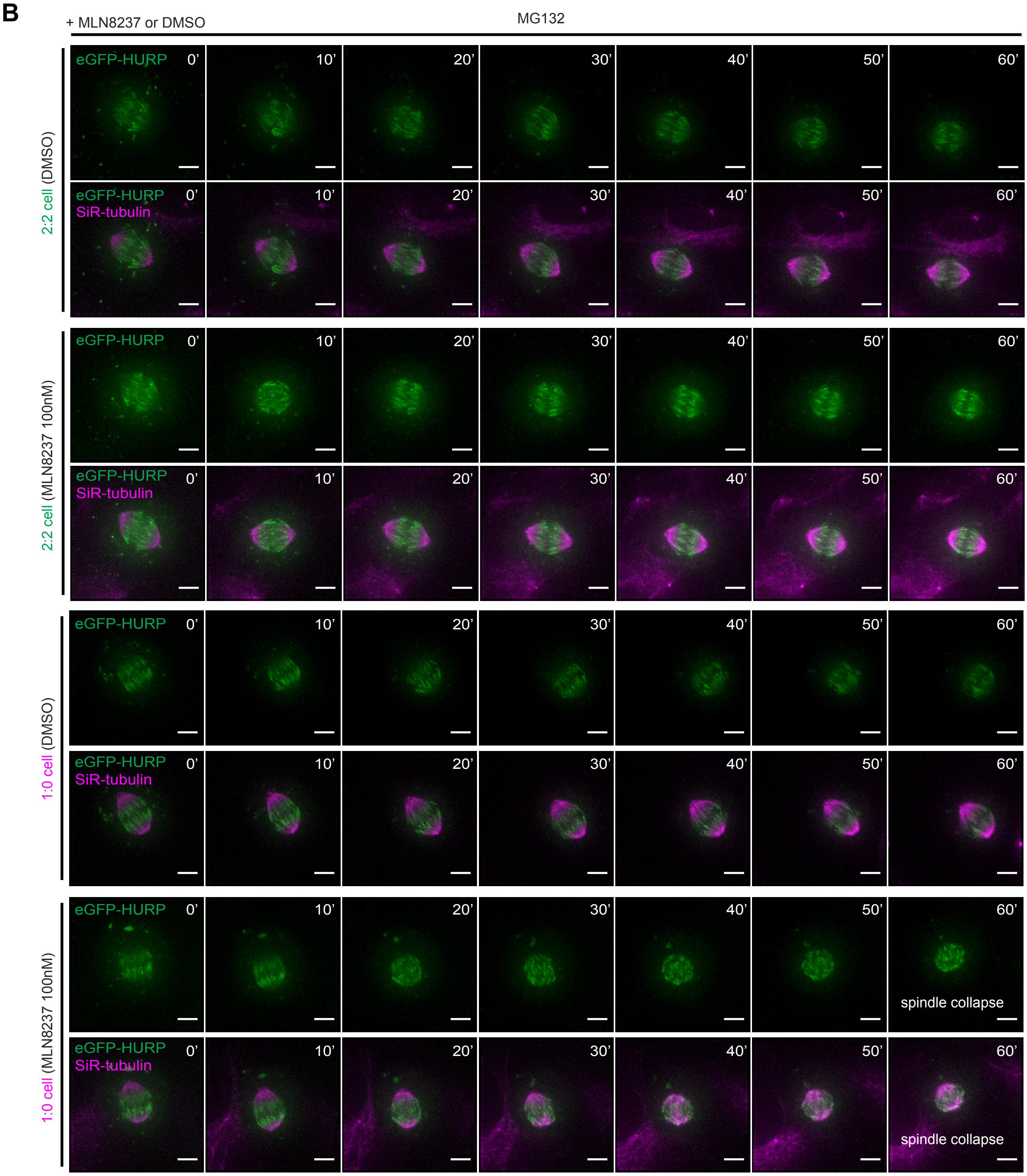

**Figure S6 (Related to Figure 6). Aurora-A inhibition does not affect HURP localization** (**A**) Immunofluorescence images of representative 2:2 (treated for 1 hour with DMSO or 100 nM of MLN8237) and 1:0 hTert-RPE1 Centrin1-GFP/GFP-CENPA cells blocked in metaphase prior fixation, stained with anti-Aurora-A-T288 antibody and DAPI. Scale bars = 5 µm. (**B**) Time-lapse images of 2:2 and 1:0 hTert-RPE1 eGFP-HURP cells incubated with 25 nM SiR-DNA, blocked in metaphase and treated either with DMSO or 100nM of MLN8237; images were taken for 1h every 10 min.

### Dudka et al., Figure S7 (Related to Figure 6)

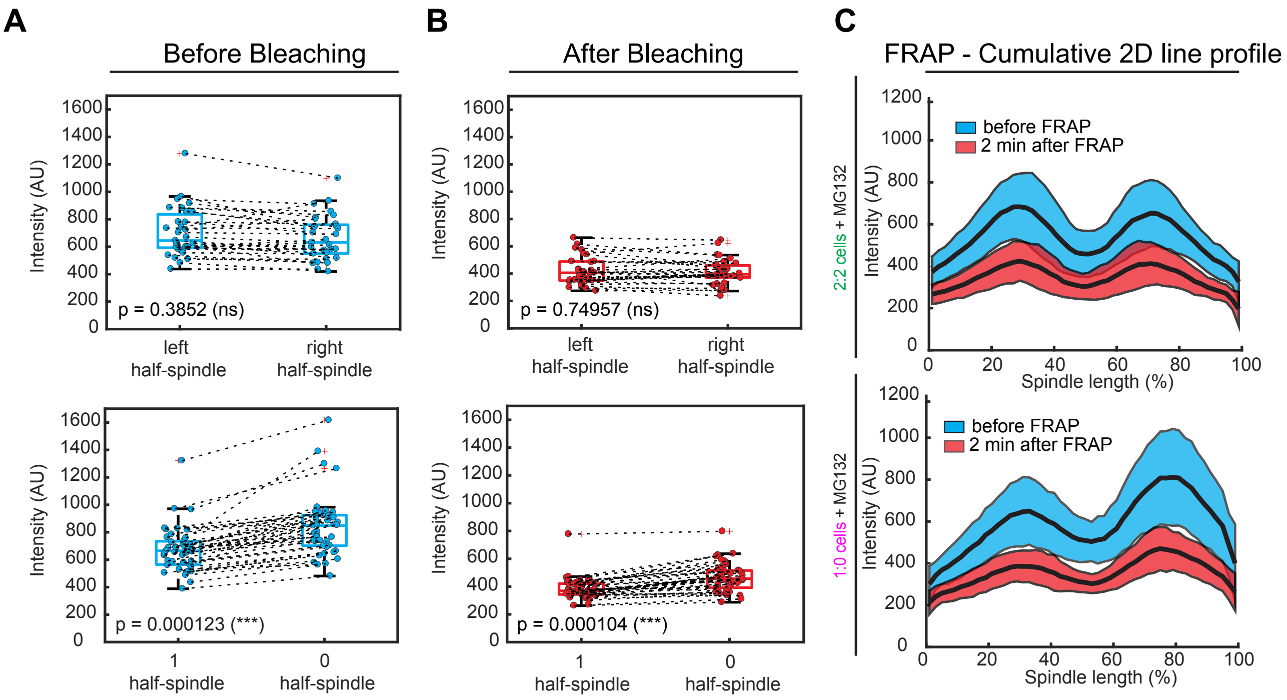

**Figure S7 (Related to Figure 6). HURP localization is linked to spindle asymmetry** (**A**) Quantification of the symmetry of eGFP-HURP signal between the opposite half-spindles measured in 2:2 (upper panel) or 1:0 (bottom) cells before bleaching; N = 3; n = 31-43 cells; left vs. right half-spindles p = 0.3852; 1 vs. 0 half-spindles p = 0.000123 in two-tailed paired t-test. (**B**) Quantification of the symmetry of eGFP-HURP signal between the opposite half-spindles measured in 2:2 (upper panel) or 1:0 (bottom) cells 2 min after bleaching; N = 3; n = 31-43 cells; left vs. right half-spindles p = 0.7496; 1 vs. 0 half-spindles p = 0.000104 in two-tailed paired t-test. (**C**) Cumulative eGFP-HURP line profiles along the spindle axis in 2:2 (upper) and 1:0 (bottom) hTert-RPE1 eGFP-HURP cells before (blue) and 2 min after (red) bleaching; thick lines – means, thin lines – s.d.; N = 3; n = 31-43 cells. *** Indicates p < 0.001, ns – not significant.
